# Human cortical pyramidal neurons: From spines to spikes via models

**DOI:** 10.1101/267898

**Authors:** Guy Eyal, Matthias B. Verhoog, Guilherme Testa-Silva, Yair Deitcher, Ruth Benavides-Piccione, Javier DeFelipe, Christiaan P.J. de Kock, Huibert D. Mansvelder, Idan Segev

## Abstract

We present the first-ever detailed models of pyramidal cells from human neocortex, including models on their excitatory synapses, dendritic spines, dendritic NMDA- and somatic/axonal- *Na*^+^ spikes that provided new insights into signal processing and computational capabilities of these principal cells. Six human layer 2 and layer 3 pyramidal cells (HL2/L3 PCs) were modeled, integrating detailed anatomical and physiological data from both fresh and post mortem tissues from human temporal cortex. The models predicted particularly large AMPA- and NMDA- conductances per synaptic contact (0.88 nS and 1.31nS, respectively) and a steep dependence of the NMDA-conductance on voltage. These estimates were based on intracellular recordings from synaptically-connected HL2/L3 pairs, combined with extra-cellular current injections and use of synaptic blockers. A large dataset of high-resolution reconstructed HL2/L3 dendritic spines provided estimates for the EPSPs at the spine head (12.7 ± 4.6 mV), spine base (9.7 ± 5.0 mV) and soma (0.3 ± 0.1 mV), and for the spine neck resistance (50 – 80 MΩ). Matching the shape and firing pattern of experimental somatic *Na*^+^-spikes provided estimates for the density of the somatic/axonal excitable membrane ion channels, predicting that 134 ± 28 simultaneously activated HL2/L3- HL2/L3 synapses are required for generating (with 50% probability) a somatic *Na*^+^ spike. Dendritic NMDA spikes were triggered in the model when 20 ± 10 excitatory spinous synapses were simultaneously activated on individual dendritic branches. The particularly large number of basal dendrites in HL2/L3 PCs and the distinctive cable elongation of their terminals imply that ~25 NMDA- spikes could be generated independently and simultaneously in these cells, as compared to ~14 in L2/3 PCs from the rat temporal cortex. These multi-sites nonlinear signals, together with the large (~30,000) excitatory synapses/cell, equip human L2/L3 PCs with enhanced computational capabilities. Our study provides the most comprehensive model of any human neuron to-date demonstrating the biophysical and computational distinctiveness of human cortical neurons.

## Introduction

Understanding the human brain is of high priority for humankind, as is manifested by the thousands of studies published every year on the various aspects of the human brain and by the large-scale projects initiated in the last decade worldwide (Amunts et al., 2016; Koch and Jones, 2016; Markram et al., 2015; Martin and Chun, 2016; Poo et al., 2016). This is a challenging task; not only because of the complexity of the brain and the technical difficulties involved, but also because ethical limitations do not allow all of the necessary datasets to be acquired directly from human brains. Consequently, most of our present knowledge of the fine structure of the brain has been obtained from experimental animals (DeFelipe, 2015). However, certain fundamental structural and behavioral aspects are unique to humans and the functional significance of the human-specific structure should be dealt with by employing a range of specific strategies. Indeed, a major goal is to improve the current technologies for the microanatomical, neurochemical and physiological analysis of the human brain by adapting methodologies that are typically used to examine the brain of experimental animals.

The use of biopsy material obtained during neurosurgical treatment for epilepsy, or following the removal of certain brain tumors, provide an excellent opportunity to study the micro-structure of the human brain, despite the fact that different medical characteristics of the patients may modify the brain tissue. The resected tissue can be immediately immersed in the fixative and therefore the ultrastructure and quality of the labeling achieved using a variety of markers for histology and immunocytochemistry is comparable to that obtained in experimental animals (Alonso-Nanclares et al., 2008; del Río and DeFelipe, 1994). Similarly, this resected human brain tissue proved to be of great value in the 1980s and 1990s to directly study the functional properties of human brain tissue *in vitro.* These studies have mostly aimed to analyze the mechanisms underlying seizures and epileptogenesis (reviewed in Avoli et al., 2005; Köhling and Avoli, 2006). Recently, there has been renewed interest in using “non-epileptic” cortical samples (removed during surgery on brain tumors) or “non-spiking” regions with normal histology (removed at a distance from the epileptic focus) in epileptic patients, as they provide an unprecedented opportunity to study human cells and local circuits, both biophysically and computationally (Eyal et al., 2016; Mohan et al., 2015; Molnár et al., 2016; Szabadics et al., 2006; Testa-Silva et al., 2014; Tian et al., 2014; Varga et al., 2015; Verhoog et al., 2013).

The other main source of tissue to study the structure of human brain is from autopsy of control individuals. In principle, this is the only source of tissue that is free of known pathology, but it is not suitable for electrophysiological studies. Another major limitation in using autopsied tissue is the post-mortem time; the longer the post-mortem time delay the larger are the alterations observed in the measurements at all levels of biological organization (genetic, molecular, biochemical, anatomical). In previous studies we have shown that post-mortem times shorter than five hours yield excellent results using fine anatomical tools like intracellular injections in fixed material or electron microscopy techniques (Benavides-Piccione et al., 2005; Blazquez-Llorca et al., 2013; Elston et al., 2001). Thus, anatomical and physiological studies of the human brain should ideally be performed by combining data from biopsies and autopsies.

Our own recent studies on human cortical neurons have shown that they are distinguished from rodent neurons in some fundamental properties. Human L2/L3 PCs are anatomically more extended and have elaborated dendritic trees (Deitcher et al., 2017; Mohan et al., 2015); have unique membrane properties (Eyal et al., 2016) and have a large number of dendritic spines/synapses per cell (DeFelipe et al., 2002; Elston et al., 2001). These neurons are capable of tracking, via their axonal spikes, very fast modulations of their dendritic inputs and their synapses recover rapidly from depression (Eyal et al., 2014; Testa-Silva et al., 2014). Still, many other biophysical properties of human pyramidal neurons remain unknown, including the magnitude, time-course and conductance composition of their synaptic inputs, and the nature of their dendritic and somatic nonlinearities. These properties are key for constructing realistic models of these cells and, based on these models, for understanding signal processing and computational capabilities of human neurons.

To extract synaptic, dendritic and computational properties of individual neurons based on diverse morphological and biophysical experiments, an overarching theoretical framework is required. Rall’s cable theory for dendrites (Rall, 1959) and his compartmental modeling approach (Rall, 1964; Segev et al., 1995) provided such theoretical framework. Indeed, in the last few decades, detailed compartmental models have been constructed for a variety of neuron types for different species and brain regions — from flies to birds, to rodents and cats; and from hippocampus to cerebellum, basal ganglia and the neocortex. These experimentally-based models provided key insights into the mechanisms governing the large dynamic repertoire of neurons and their computational and plastic capabilities (reviews in (Herz et al., 2006; Koch and Segev, 2000; Major et al., 2013; Stuart et al., 2016)). But to what extent do these neuron models in non-human mammals (e.g., rodents) provide insights into the biophysics and computational capabilities of human neurons? In our recent work (Eyal et al., 2016) we constructed passive compartmental models of HL2/L3 PCs to study their membrane properties. To our surprise, we found that the specific membrane capacitance (*C*_*m*_) of human neurons is distinctive; ~0.5 μF/cm^2^, half of the commonly accepted “universal” value (~1 μF/cm^2^) for biological membranes. The low *C*_*m*_ has important functional implications for signal processing in these cells. This initial surprise has led us to perform hereby a more comprehensive modeling study of HL2/L3 PCs

In the present work, we integrated a wide range of anatomical and physiological data on six HL2/L3 PCs, including the fine anatomy of dendritic spines, which provided the most comprehensive models of human cortical pyramidal neurons to date. We extended our existing passive models of HL2/L3 PCs in order to estimate their synaptic properties, in particular the NMDAR-kinetics, and to estimate the properties of the ion channels underlying the somatic/axonal spiking mechanisms. Based on these parameters, we predicted the conditions for the generation of NMDA spikes in individual dendritic branches receiving excitatory spinous synapses, as well as the number of excitatory synapses required to initiate axo-somatic (output) *Na*^+^ spikes. Our models show that human L2/L3 PCs have the capacity to generate tens of independent dendritic NMDA spikes (supporting local nonlinear dendritic computations). We further show that, despite the extended dendritic cable structure of human L2/L3 PCs, a relatively small number (~135) of synchronously activated excitatory axo-spinous synapses is required to generate an axonal output spike. We concluded that human L2/L3 pyramidal cells expand the computational/memory capacity of similar cells, e.g., in rodents, because of their increased numbers of local dendritic nonlinear subunits, increased excitability due to low *Cm* value and their large number (~30,000) of dendritic spines/excitatory synapses.

## Material and methods

### Experimental data

#### Human brain slice preparation

All procedures on human tissue were performed with the approval of the Medical Ethical Committee (METc) of the VU University Medical Centre (VUmc), with written informed consent by patients involved to use brain tissue removed for treatment of their disease for scientific research, and in accordance with Dutch license procedures and the declaration of Helsinki (VUmc METc approval ‘kenmerk 2012/362’). After resection, the neocortical tissue was placed within 30 seconds in ice-cold artificial cerebrospinal fluid (aCSF) slicing solution which contained in (mM): 110 choline chloride, 26 NaHCO3, 10 D-glucose, 11.6 sodium ascorbate, 7 MgCl2, 3.1 sodium pyruvate, 2.5 KCl, 1.25 NaH2PO4, and 0.5 CaCl2 – 300mOsm, saturated with carbogen gas (95% O2/ 5% CO2) and transported to the neurophysiology laboratory located 500 meters from the operating room. The transition time between resection of the tissue and the start of preparing slices was less than 15 minutes. Neocortical slices (350-400 μm thickness) were prepared in ice-cold slicing solution, and were then transferred to holding chambers filled with aCSF containing (in mM): 126 NaCl; 3 KCl; 1 NaH2PO4; 1 MgSO4; 2 CaCl2; 26 NaHCO3; 10 glucose – 300mOsm, bubbled with carbogen gas (95% O2/ 5% CO2). Here, slices were stored for 20 minutes at 34°C, and for at least 30 minutes at room temperature before recording.

#### 3D reconstructions of dendritic arbors and dendritic spines of HL2/L3 pyramidal cells

Six dendritic morphologies of L2/L3 from human temporal cortex, residing at the depths of 675-1204 μm below the pia were used in this study. These are the same morphologies used in (Eyal et al., 2016), taken from (Mohan et al., 2015). These neurons were recorded and then labeled using biocytin as a marker that was revealed with DAB using the chromogen 3,3-diaminobenzidine tetrahydrochloride and the avidin–biotin–peroxidase method.

For data on dendritic spines we used L3 pyramidal cells that were reconstructed with a confocal microscope after intracellular injection with Lucifer Yellow in the temporal cortex, corresponding to Brodmann’s area 20 (Brodmann, 2007). These neurons were obtained from two human males (aged 40 and 85) obtained at autopsy (2–3 h post-mortem) following traffic accidents. This human material has been used in a previous study (Benavides-Piccione et al., 2013). We used data on dendritic spines from the post-mortem tissue instead of directly measuring dendritic spines from the recorded cells because confocal microscopy of Lucifer Yellow-labeled neurons is more appropriate to analyze the shape and the size of dendritic spines than light microscopy of biocytin-labeled neurons. The complete morphology of over 8300 dendritic spines was reconstructed in 3D as in (Benavides-Piccione et al., 2013). Briefly, spines and dendritic branches were imaged using a Leica TCS 4D confocal scanning laser attached to a Leitz DMIRB fluorescence microscope. Consecutive stacks of images (z-step of 0.28 μm) were acquired using a 0.075×0.075×0.28 μm3 voxel size (Leica Objective Plan-Apochromat 63x/1.30 NA glycerol DIC M27) to capture the full dendritic depths, lengths, and widths of the dendrites (**Figure 2**). Dendritic spine structure was analyzed in 20 basal dendrites and 16 main apical dendrites (10 basal dendrites and 8 main apical dendrites per case), using Imaris 6.4.0 (Bitplane AG, Zurich, Switzerland). Correction factors used in other studies in which dendritic spines were quantified in opaque material (e.g. Golgi method or biocytin/DAB material) were not used in the present study as the fluorescent labeling and the high-power reconstruction allowed the visualization of dendritic spines that protrude from the underside of dendrites. Since confocal stacks of images intrinsically result in a z-dimension distension, a correction factor of 0.84 was applied to that dimension. This factor was calculated using a 4.2 μm Tetraspeck Fluorescent microsphere (Molecular Probes) under the same parameters used for the acquisition of dendritic stacks. No optical deconvolution was used for spine reconstruction. Finally, dendritic spines with a head diameter below the resolution limit were not considered in the present study. However, for the purposes of modeling, this factor is probably insignificant since, in a previous electron microscopic study, it was found that “thin” dendritic spines (i.e., those that lacked clear heads, resembling the filopodia found at earlier developmental stages) are non-synaptic (Arellano et al., 2007).

**Figure 1.**
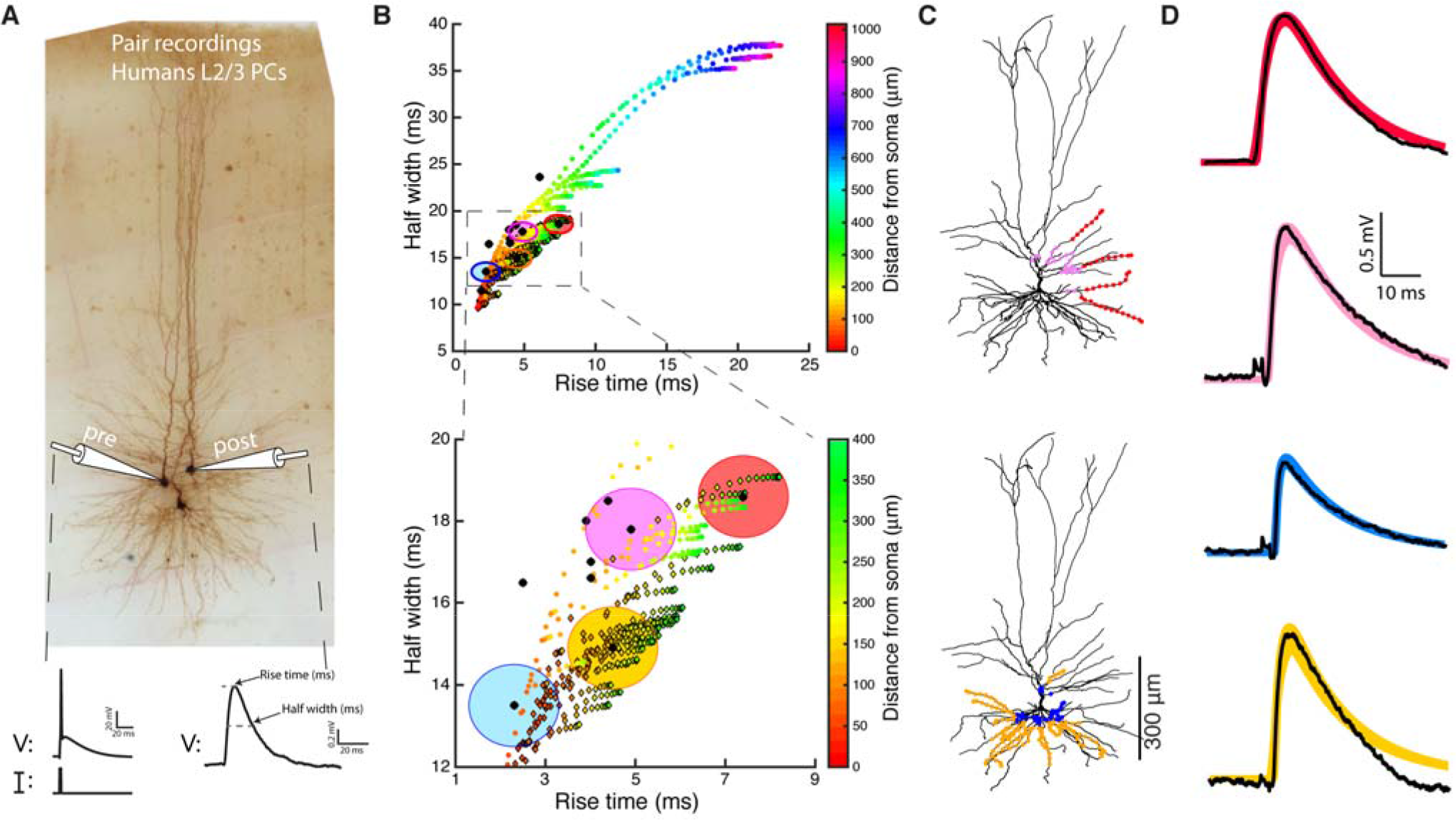
Model predicts that HL2/L3 - HL2/L3 excitatory synapses are formed at proximal dendritic sites. **(A)** Pair recording from HL2/L3 PCs. A presynaptic spike was initiated in a cell (lower left trace) and the post synaptic EPSP was measured in another cell (lower right trace). The shape index of this EPSP is defined by its rise time and half width (bottom right). **(B)** Top. Theoretical shape-index curve for the modeled cell shown in **(C)**, as a function of distance from the soma. Colors code for the physical distance from the soma; color circles for apical inputs and color diamonds for basal inputs. Bottom. Zoom-in into the square demarcated at the top frame. Black circles are from ten experimental somatic EPSPs. The large filled color circles with radius of 1 ms are centered around the loci of the respective four experimental EPSPs shown in **(D)**. **(C)** Modeled cell used in **(B)**, with dots depicting the predicted synaptic locations that give rise to somatic EPSPs whose shape indices fall within the corresponding large colored circles in (**B)**. E.g., red points are all synaptic contacts that yield rise-time and half-width that are within the red circle in **(B)**. **(D)** Four experimental EPSPs (black traces) from four connected pairs of HL2/L3-HL2/L3 pyramidal cells and the theoretical EPSPs (thick color) corresponding to the respective color dots in **(C)**. The peak synaptic conductance, for each of the putative dendritic synapse, was obtained via fitting the theoretical to the experimental transients (see text and **Table S1**). The recordings in **(A)** were taken from a pair of cells that were not reconstructed, and the HL2/L3 morphologies are shown here only for the illustration of the method (See **Figure S1**).

For modeling of typical dendritic spines, which establish at least one synaptic contact, we measured the spine neck length, spine neck diameter and spine head area of a selection of 150 dendritic spines that clearly showed a spine head. Only, spines showing a clear head whose morphology could be captured using a single surface of a particular intensity threshold were included in the study. The spine neck length and spine neck diameter were manually marked in each selected dendritic spine from the point of insertion in the dendritic shaft to the spine head, while rotating the image in 3D (**Figure 2**).

#### Electrophysiology (acute living slices)

Whole-cell, patch clamp electrophysiology recordings were made from human L2/L3 pyramidal neurons as described previously (Testa-Silva et al., 2014; Verhoog et al., 2013). Whole-cell recordings were made using uncoated, standard borosilicate glass pipettes. Recordings were made using multiclamp 700B amplifiers (molecular devices) and were digitized using Axon Instruments Digidata 1440A. Recording aCSF matched the solution of the aCSF in which slices were stored. The recording temperature was 32–35°C. Internal solutions were (in mM): 110 Kgluconate; 10 KCl; 10 HEPES; 10 K2Phosphocreatine; 4 ATP-Mg; 0.4 GTP, biocytin 5 mg/ml, pH adjusted with KOH to 7.3 (290–300 mOsm).

#### EPSPs from paired recordings

Unitary EPSPs were measured from ten connected pairs of L2/L3 human pyramidal neurons (Testa-Silva et al., 2014). These experimental EPSPs came from a different set of neurons than those used in (Eyal et al., 2016). We therefore used experimental EPSPs measured in HL2/L3 PCs that exhibit cable properties (input resistance, membrane time constant) similar to our neuron models. The membrane time constant for the post synaptic cell from which the EPSP was recorded was determined by “peeling” the tail of the EPSP (Rall, 1969). Only cells with time constant of 17±5 ms were considered. Protocols for experiments are available in (Testa-Silva et al., 2014).

#### Extracellular stimulation and NMDA data

EPSPs were evoked every seven seconds by extracellular stimulation using two bipolar stimulating electrodes in glass pipettes loaded with extracellular/recording aCSF that were positioned extracellularly, 100-150 μm away from the soma and approximately 50 μm lateral to the cell’s apical dendrite. Duration (50 μsec) and amplitude (~40 μA, range 30-70 μA) of extracellular stimulation were controlled by Isoflex stimulators (A.M.P.I.). For each cell, two extracellular stimuli were applied at two different loci with respect to the cell body of the post-synaptic cell (6 experimental EPSPs). Seventeen EPSPs were evoked and averaged before the use of blockers. Then, NMDA-receptor mediated EPSPs were isolated by blocking AMPA and kainite receptors with NBQX (1μM, bath-applied, solved in recording aCSF) and blocking GABA_A_ with Bicuculline or GABAzine (10 μM, bath-applied, solved in recording aCSF). For one of these cells (**Figure 4B)** we already had a passive model from (Eyal et al, 2016). The model was based on brief current injections into the cell as well as its 3D morphology (Eyal et al., 2016, Figure 1c5). We used this model when fitting the NMDA-receptor kinetics (**Figure 4C**, see below for more details).

**Figure 2.**
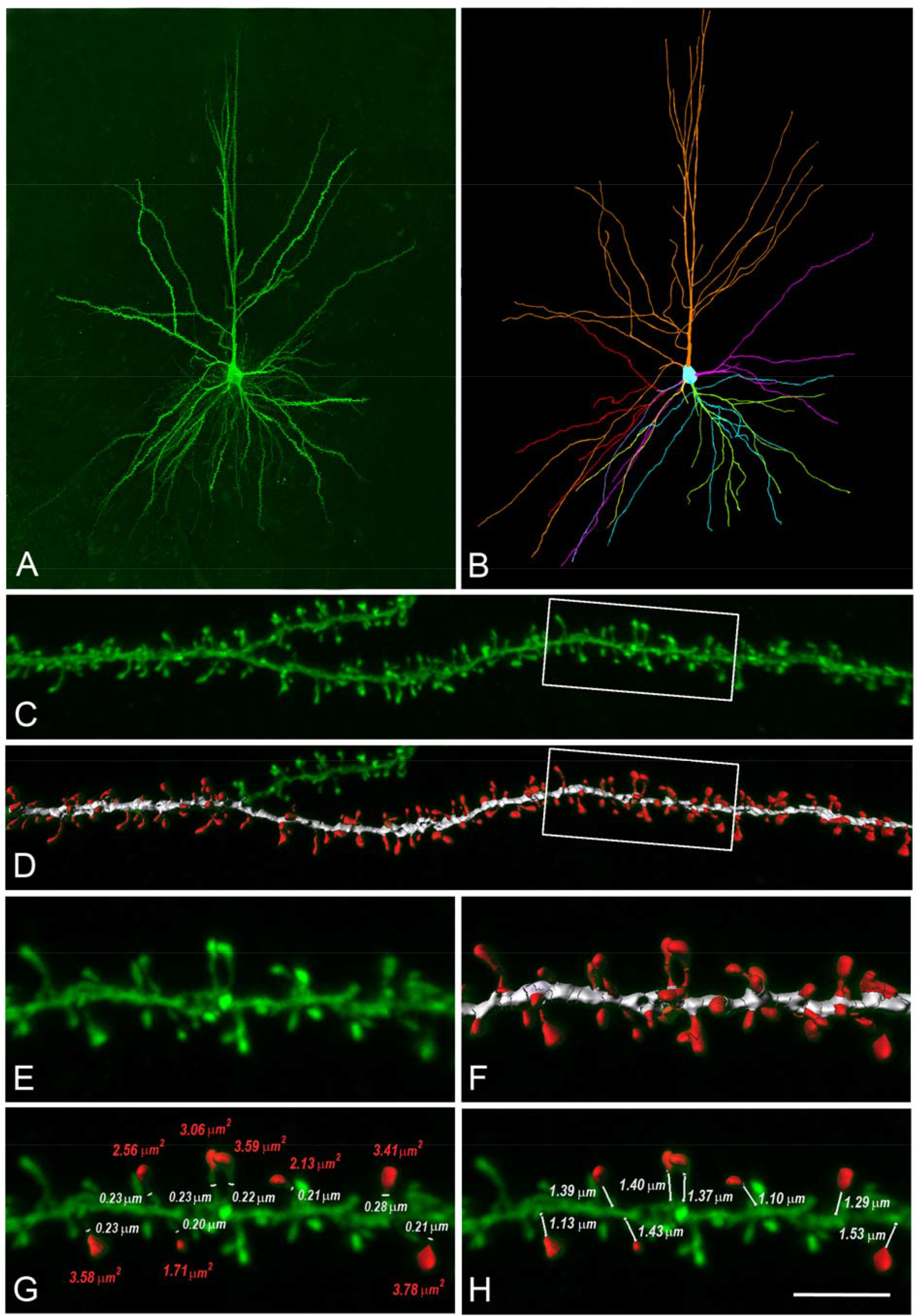
Reconstructions of human L3 dendritic spines. **(A)** Confocal microscopy image z projection of an intracellular injected layer 3 pyramidal neuron of the human temporal cortex obtained at autopsy. **(B)** 3D reconstruction of the complete morphology of the cell shown in **(A)**. Orange represents the apical dendritic arbor whereas other colors represent the basal dendritic arborization. **(C)** Confocal microscopy image showing a horizontally projecting labeled basal dendrite. **(D)** To reconstruct the complete morphology of dendritic spines (red), different intensity thresholds were created and then a particular threshold was selected for each spine to constitute a solid surface that exactly matched its contour. The dendritic shaft (white) was 3D reconstructed by selecting a particular threshold that represented a solid surface that matched the contour of the dendritic shaft along the length of the dendrite. **(E, F)** Higher magnification images of the dendritic segment indicated in boxed areas in **(C)** and **(D)**. **(G)** For a selection of spines which showed clear heads, a particular solid surface that matched the contour of the spine head was created (red). The neck diameter was manually marked (white). Spine head area and neck diameter measurements are indicated in red and white numbers, respectively. **(H)** The neck length was manually marked from the point of insertion in the dendritic shaft to the spine head. Neck length measurements are indicated in white numbers. Scale bar (in **(H)**): 110 μm in **(A)**, **(B)**, 10 in **(C, D)** and 4.5 μm in **(E–H)**.

#### Train of somatic action potentials

A step of depolarizing current of 1 sec long was injected to the somata of the six HL2/L3 pyramidal cells that are the focus of this paper. Current amplitudes were adjusted so that each cell fired at about ~10 Hz (examples are shown in **Figure 7A**). This stimulus was repeated 10 times or more per cell, providing sufficient statistics to use the multiple objective optimization (MOO) to develop channel-based models for these spikes (see below). All traces were sampled at 100 - 250 kHz and low-passed filtered at 14 - 15 kHz.

#### I-F curves of HL2/L3 PCs

We computed the I-F curve for 25 additional HL2/L3 PCs (I-F curves for the six modeled cells were not available) as part of our modeling efforts to match experimental results to model performance using the MOO algorithm (see below). These I-F curves were computed from the spike trains following 1 sec depolarizing step current of different supra-threshold amplitudes. Nearby points in the I-F curve were connected by linear lines; individual I-F curves were then normalized by the input current that lead to 10 Hz firing rate (see also (Hay et al., 2011)). The 25 I-F curves and their mean are shown in **Figure 7B** (grey and black traces respectively).

### Modeling

#### Simulations

Simulations were performed using NEURON 7.4 (Carnevale and Hines, 2006) running on a grid of 60 Sun 4100 AMD 64-bit Opteron dual core (240 cores in total), running Linux 2.6, or on grid of 40 Intel(R) Xeon(R) CPU E5-2670 with 16 cores per node (640 cores in total), running Redhat 6.6.

#### Passive neuron models

In (Eyal et al., 2016) we constructed detailed passive compartmental models for six 3D reconstructed L2/L3 pyramidal neurons from human temporal cortex. The three passive parameters (*C*_*m*_, *R*_*a*_, *R*_*m*_) in these models were optimized for each modeled cell such that the theoretical transients following brief/small current steps generated by the model closely fit the corresponding experimental transient. Details could be found in (Eyal et al. 2016). One key result from this study was that in human neurons, *C*_*m*_ is half (0.5 μF/cm^2^) than the “universal” value of 1 μF/cm^2^. In the present work, we use these six models as the passive skeleton onto which experimentally-constrained synaptic and membrane nonlinearities were added.

Membrane area of the dendritic spines, which are abundant in human pyramidal cells, was incorporated globally into the 3D reconstructed dendritic models using the factor *F*,

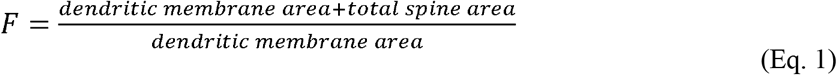

The incorporation of dendritic spines into a particular dendritic branch was implemented by multiplying *C*_*m*_ by *F* and dividing *R*_*m*_ by *F* as described previously (Rapp et al., 1992). Spine and shaft areas were computed using reconstructions of 3D images from confocal microscopy of samples from two post mortem brains (Benavides-Piccione et al., 2013, see **Figure 2** below). This resulted with an *F* value of 1.9. Spine membrane area was incorporated into the modeled neuron only in dendritic segments that are at a distance of at least 60 μm from the soma, due to the low density of spines in more proximal branches (Benavides-Piccione et al., 2013). More details can be found in (Eyal et al., 2016).

#### Model for dendritic spines

Individual dendritic spines receiving excitatory synaptic input were modeled in details (**Figures 3**–**6** and **8**) using two compartments per spine; one for the spine neck and one for the spine head. The spine neck was modeled using a cylinder of length 1.35 μm and diameter of 0.25 μm, whereas the spine head was modeled as an isopotential compartment with a total area of 2.8 μm^2^. The dimensions for these compartments were based on measurements from 3D reconstructed dendritic spines of human temporal cortex L3 pyramidal cells (see the “3D reconstructions of dendritic arbors and dendritic spines of HL2/L3 pyramidal cells” section above). The passive parameters (*C*_*m*_, *R*_*m*_, *R*_*a*_) of the spine were similar to those of the dendrites. This spine model led to a spine neck resistance of 50-80 MΩ.

#### Synaptic inputs

We simulated AMPA-based and NMDA-based synaptic currents as follows,

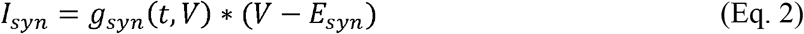

where *g*_*syn*_ is the synaptic conductance change and is the reversal potential for the synaptic current.*E*_*syn*_ was 0 mV for both the AMPA and the NMDA currents.

The synaptic conductance was modeled for both AMPA and the NMDA components, using two state kinetic scheme synapses – with rise time (τ_*rise*_) and decay time (τ_*decay*_) constants:

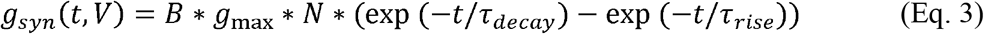

Here *g*_*max*_ is the peak conductance and *N* is a normalization factor given by

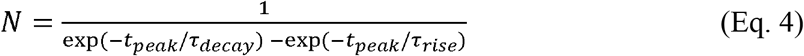

and *t*_*peak*_ (time of the peak conductance) is calculated as:

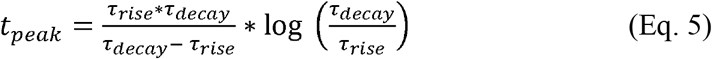

AMPA kinetics was kept constant throughout this study and only its peak conductance was fitted for the various cases studied. For modeling AMPA-based conductance, *B* was set to 1 (voltage-independent conductance). Standard values for τ_*rise*_ and τ_*decay*_ were 0.3 ms and 1.8 ms respectively. We tried other values but this led to a poorer fit between the theoretical and experimental EPSPs shown in **Figure 1D**.

NMDA conductance is voltage dependent. In this work, B was defined using the equation as in (Jahr and Stevens, 1990):

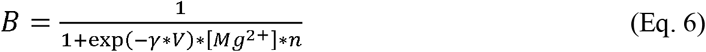

Mg^2+^ concentration was 1mM in the model and other parameters were computed so that they best fit the experimental results (see **Figure 4**, and section “Modeling the NMDA-based current to fit the experimental results”).

#### Shape index curves for estimating the dendritic loci of HL2/L3-HL2/L3 synaptic connections

Experimental EPSPs were measured via patch recordings from ten synaptically-connected L2/L3-L2/L3 neuron pairs (Testa-Silva et al., 2010). The EPSPs were recorded from the soma of the post synaptic neuron, following the activation of a single spike at its presynaptic L2/L3 neuron. These ten connected cell pairs were not reconstructed in 3D. We therefore selected one 3D reconstructed L2/L3 neuron as our prototypical neuron (see **Figure 1C** and also Figure 1A in (Eyal et al., 2016)) and used this model to construct a “shape index curve” (EPSP rise-time versus half width) as in (Rall et al., 1967) in order to estimate the putative location of the excitatory synapses that gave rise to the experimental EPSPs. Although it would be preferable to model each one of the post-synaptic cell from which the EPSPs were recorded from, we believe that the morpho-electrotonic variance among HL2/L3 PCs is sufficiently small so that our conclusions, using a prototypical neuron for constructing the “shape index curve”, for characterizing the dendritic “territory” of HL2/L3 - HL2/L3 synaptic connections and the range of their conductance values is valid.

For simplicity, we started by assuming that HL2/L3 – HL2/L3 connection is formed by a single synapse. The theoretical shape index curve was calculated whereby each of the model electric compartments was activated by a single AMPA-synapse (see above). Then, we superimposed the experimental EPSP half-width versus rise time on this theoretical curve (black dots in **Figure 1B**). We defined an electric compartment in the model to be a putative synaptic contact if its shape index value was inside a circle with a diameter of 1 ms around the experimental EPSPs shape index (four of these circles are shown in **Figure 1B** bottom). Then we used NEURON’s PRAXIS optimizer (Brent, 1976; Carnevale and Hines, 2006) to compute the peak AMPA conductance (*g*_*AMPA*_) for each putative synapse that resulted with the best fit to the experimental EPSP (**Figure 1D**). The estimated synaptic peak conductance was calculated as the average peak conductance over all putative locations.

Next, we assumed that five synaptic contacts are formed per HL2/L3 – HL2/L3 connection, similar to the average value for L2/3 and L5 PCs in rodents (Feldmeyer et al., 2002, 2006; Markram et al., 2015). For each group of putative synapses obtained above, we randomly selected five synapses, from the suggested putative locations when assuming a single contact per axon, and activated them simultaneously. We then ran the optimization procedure to obtain the peak conductance/contact. This process was repeated 100 times and the mean AMPA conductance value/contact was estimated as shown in **Table S1**.

The above estimations were performed assuming that, for a single connection, only AMPA conductance is activated. This is justified as the activation of a single synaptic connection generates local voltage that is too small (8.6 ± 6.5 mV) to activate a significant NMDAR current (**Figure 5**). This allowed us at a later point to build a model for the NMDA-kinetics and conductance amplitude that is based on the results from the earlier part (see below, and **Figure 4**). We validated that the addition of NMDA conductances to the modeled synapses does not change significantly the results of **Figure 1** and **Table S1** in **Figure S2**.

#### Modeling the NMDA-based current to fit the experimental results

The process of fitting the NMDA model kinetics to the experimental results was based on having both the morphology of the post-synaptic cell (**Figure 4B**), the somatic voltage transients (Eyal et al., 2016, Figure 1d5) and the composite EPSPs recorded from the soma following extracellular simulation with and without AMPA blockers, all from the same cell (**Figure 4C**). This allowed us to use the passive model of this cell from (Eyal et al., 2016) and, on top of it, add AMPA- and NMDA- based currents such that the model fits the properties of this cell extracellular-generated somatic EPSPs (**Figure 4**).

The optimization of the synaptic parameters that fit the experimentally-recorded composite EPSP was achieved as follows. First, between 15 to 30 synapses were randomly distributed in a restricted part of the apical tree (near the location of the extracellular electrode); each of the synapses having both NMDA and AMPA conductance. For the optimization, the NMDA peak conductance and its kinetics (τ_*rise*_ and τ_*decay*_ in **Eq. (5)** and the “steepness” factor γ in **Eq. (6)**), as well as the AMPA peak conductance were free parameters. The parameter *n* in **Eq. (6)** remained constant in this work with a value of 0.28 1/mM as found by (Jahr and Stevens, 1990) for an extracellular solution with 1mM magnesium concentration. We also tried other available models for the NMDA kinetics (Rhodes, 2006, Sarid et al., 2007), but those led to a poorer fit of the data. The optimization was achieved by first fitting the model response to the experimental EPSP in the presence of AMPA blockers. In the experiments, a competitive antagonist was used to block the AMPA receptors, therefore in the simulations we allowed the model to include a small AMPA conductance (see below). This implied that, even in the presence of an AMPA blocker, a strong extracellular stimulus could activate a small APMA current which, in turn, could help activate the NMDA-current, as indeed was found experimentally.

For these simulations, 60,000 seeds were used to select the number of synapses and their dendritic location; each one of them was optimized using the PRAXIS algorithm in NEURON (Brent, 1976; Carnevale and Hines, 2006). The top 20,000 models were chosen for the next step. In this stage, we tried to fit the model with the experimental EPSP recorded without any blockers. In each model, using the result of the first stage of the fit, the synaptic locations and NMDA conductance and kinetics were set, with the only free parameter at this stage being peak AMPA conductance. The peak AMPA conductance was constrained to be at least five times larger than its peak conductance on the first stage (the blocked case). Models were sorted according to their sum root mean square distance, RMSD, with respect to the two target experimental EPSPs. Different locations of the synapses and other optimization methods were also attempted; eventually the procedure described above resulted with a set of 100 models that best fit the experimental EPSPs (see **Figures 4** and **S3 and Table S2**).

We selected one of the best five models resulting from the above procedure for the rest of this work. Out of best five, we chose the model with the parameters that were the closest to the mean of the best 100 models (best typical). The corresponding kinetics of the model for the NMDA conductance were: τ_*rise*_ = 8.02 ms, τ_*decay*_ = 34.99 ms, 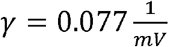. The conductance values per contact were as follows: *g*_*NMDA*_ = 1.31 *nS*, *g*_*AMPA*_ = 0.73 *nS* and 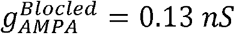 The values of τ_*rise*_, τ_*decay*_ and γ were within the range found in rodents (Sarid et al., 2007), and used by many other studies (e.g., Rhodes, 2006). It is important to note that all the best 30 models had γ values larger than 0.075 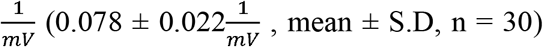 implying a relatively steep dependency of the NMDA- current on voltage as compared to rodents (see below).

#### NMDA spikes

In **Figure 5** we modeled an NMDA spike that originated from synapses distributed along a 20 μm stretch of a dendritic segment. Synapses were activated on the spine head membrane and the resultant voltage was recorded in the spine head, the dendritic shaft and the soma. In **Figure 6** we computed the maximal number of independent NMDA spikes (maximal number of independent nonlinear dendritic subunits) that could be generated by the modeled cell. Synapses were located in the distal dendritic terminals and the number of synapses in each terminal was the minimal number required for generating a local NMDA spike, defined as such when the local voltage is larger than −40 mV and lasts for at least 20 ms. We found that this threshold is a good criterion for distinguishing between a brief and strong AMPA-based EPSPs and a prolonged NMDA+AMPA based spikes (also known as plateau potentials). These NMDA spikes usually reached maximal local depolarization that is close to the reversal potential of NMDA current (0 mV in the present model). NMDA spikes were defined as independent from each other if the peak voltage in the branch-point connecting two activated terminals was below −40 mV. The locations and the number of the maximal clusters that generated independent NMDA spikes were chosen both manually (adding clusters of spinous inputs in the most distal locations in electrotonic terms) as well as with recursive algorithms. At the end of these procedures we could assess the maximal number of independent NMDA spike in any given neuron model.

#### Fitting trains of somatic Na^+^ spikes

The fit of the fully active axon-soma models was achieved using the multiple objective optimization (MOO) method as in (Druckmann et al., 2007; Hay et al., 2011). For obtaining a fit between model and experimental spikes, as shown in **Figure 7**, at least ten repetitions of identical depolarizing step current injections leading to about 10 Hz firing rate were experimentally recorded in the cells that were later 3D reconstructed and modeled in detailed. I-F curves were also recorded for an additional set of 25 HL2/L3 PCs (see above). The MOO procedure is based on deconstructing the spike properties to a set of features (see below) and using their experimental mean and standard deviation (from all experimental traces) to find the peak conductance of a set of predefined modeled excitable ion channels that best fit the experimental spike firing. The features for the MOO algorithm used in this study were: 1. Voltage base – the mean membrane voltage before the stimulus. 2. Steady state voltage – the mean voltage at the end of the stimulus. 3. Mean frequency – firing rate mean frequency. 4. Time to first spike – the time in ms between the stimulus onset and the peak of the first spike. 5. Burst Inter Spike Interval – the length (in ms) of inter-spike interval (ISI) between the first two spikes. 6. Inter-spike interval coefficient of variance – defined as ISI_mean_/ISI_SD_. 7. Adaptation index – normalized average difference of two consecutive ISIs. 8. AP height – average peak voltage of the spikes. 9. AP begin voltage – the voltage at the beginning of the spike, defined as the membrane voltage where dV/dt crosses 20 mV/ms. 10. After Hyper-Polarization Depth – the minimum voltage between spikes. 11. After Hyper-Polarization time from peak – the duration it takes to reach maximal hyperpolarization following the peak of the spike. 12. Spike half width – the width of the spike in its half height. 13. The mean firing rate of the normalized I-F curve of 25 HL2/L3 PCs (see above), for an input that is 75% compared with the input current with the mean frequency in Feature #3. **(Figure 7B).** For example, 10 repetitions of 700 pA to the soma of 0603_cell03 (blue morphology in **Figure 7**, second column in **Table S3**), resulted in a 10.38 ± 0.31 Hz firing rate. The normalized input for the 25 cells that lead to 10.38 Hz is 1.02 (where 1.0 is the input results with 10 Hz). 75% of 1.02 is 0.765. The mean frequency corresponding to 0.765 normalized input is 3.43 Hz ± 2.03, therefore this value is feature #13 for this cell. 14-17. As in feature #13, but for inputs of 125%, 150%, 200% and 300% respectively. The values (mean ± SD) for the different features of the six cells is provided in **Table S3**. Features were extracted using the eFEL library in python. Equations for the various features could be found in https://github.com/BlueBrain/efel.

A short axon of 60 μm was added to the modeled cells, similar to (Hay et al., 2013; Markram et al., 2015). The parameter set for the optimization consisted of 29 free parameters: the maximal conductance of nine ion channels, three kinetics parameters for the sodium, two parameters for the intracellular Ca^2+^ concentration, all were fitted both for the soma (14 parameters) and for the axon (14 parameters). The last parameter was the reversal potential of the leak current (same value for all the compartments in the model). The full list of parameters and their values for the six models is provided in **Table S4**. Since we still lack dendritic recordings from human cells, it might not be possible to constrain dendritic parameters based only on somatic recordings (Shen et al., 1999) and therefore we decided to include active parameters only in the soma and in the axon.

Optimization of the spiking activity for HL2/L3 PCs was achieved using MOO combined with an evolutionary algorithm under the Optimization-Framework of the Blue Brain Project (Markram et al., 2015). The optimization algorithm is explained in details in (Druckmann et al., 2007; Hay et al., 2011). Briefly the evolutionary process starts with 1000 random models (random set of parameters). In each generation, only the models that are the most successful are selected to pass over to the next generation (see below). In each generation, new models are generated using mutations from the successful models of the previous generation. The proximity of the models to the objectives (target features) is defined by the distance, in standard deviations, between the model feature and its respective experimental feature. Models are defined as successful if they are not dominated by any other models. i.e. there is no single model that is more successful in all the objectives. The optimization stopped after 500 generations and, for the purposes of this work, we took one model that survived through the last generation and had a small distance in all the objectives.

#### Number of activated dendritic spines synapses per somatic spike

Synapses with both AMPA- and NMDA-based conductances, as in **Figure 4**, were randomly distributed over dendritic spines (**Figure 8C**). The spatial distribution of synapses was either random or clustered-random. In the first case, the location of each synapse was chosen from a uniform distribution in the modeled dendritic tree. In the second case, the locations of a group of spatially clustered synapses was uniformly distributed over the dendritic tree; each cluster included 20 synapses located in the same dendritic branch all located within 20 μm from each other. The synapses were activated synchronously and the simulated somatic voltage was recorded. A somatic Na^+^ spike was marked as such when the somatic voltage crossed a voltage threshold of 0 mV. Each experiment (different total number of synapses/clusters in each model) was repeated 1000 times with different seeds. To estimate the number of synapses required to generate a somatic/axonal Na^+^ spike in 50% of the cases (**Figure 8B**), a linear extrapolation was used for the clustered case.

#### Rat L2/3 PC models

For the comparison between the new models of human L2/L3 neurons and existing models of rodents we used three models of L2/3 pyramidal cells from the rat neocortex. Two cells were obtained from the barrel cortex (Sarid et al., 2013, models 110602A and 280503A), and one model from the somatosensory cortex (Markram et al., 2015, model L23_PC_cADpyr229_5). All three models were used to compute the number of independent NMDA spikes that could be generated in a rat pyramidal neuron. The neuron model taken from the rat somatosensory cortex was used for the estimation of the number of excitatory synapses that were required to generate somatic Na^+^ spike in a rat L2/3 PC.

## Results

### Synaptic connections between HL2/L3-HL2/L3 PCs are proximal and powerful

We studied the properties of synaptic connections between human L2/L3 PCs using the shape index curve (EPSP half width versus its rise time) as proposed by Rall (Rall, 1967; Rall et al., 1967, see **Methods**). We used the time-course and magnitude of somatic EPSPs recorded experimentally from ten synaptically-connected pairs of HL2/L3 PCs (Testa Silva 2014, and **Figure 1A**), combined with a detailed model of a HL2/L3 PC (**Figure 1C**). The shape index values for these ten experimental EPSPs are shown by black dots in **Figure 1B**. The modeled synapses shape indices are shown as colored circles (apical synapses) and colored diamonds (basal synapses) in **Figure 1B**; colors denote the physical distance of the synapse from the soma. Comparing the experimental and the theoretical results shows that the former has a relatively brief rise time and a narrow half-width. A magnification of the initial part of the theoretical shape index curve is depicted in **Figure 1B** Bottom. This result clearly indicates that the experimental recorded EPSPs originated from relatively proximal synaptic contacts.

Based on the theoretical shape index curve we found the putative dendritic location of the synapses that give rise to each of the experimental EPSP. This location was computed so that the theoretical shape index of the putative synapses falls within a radius of one millisecond from the respective experimental shape index (large colored circles in **Figure 1B**). The locations of putative synapses for four cases are indicated by the corresponding colored dots superimposed on the dendrites of the modeled HL2/L3 PC shown in **Figure 1C**. For example, the experimental EPSP in Figure 1D **top trace**, with its rightmost shape index in **Figure 1B** (black dot surrounded by the red circle), may arise from any of the synapses whose shape index falls within the red circle. Our computations show that these putative synapses could be located on two basal dendrites and two proximal oblique dendrites, as illustrated by the red dots superimposed on the modeled cell (**Figure 1C**, top). Other three experimental EPSPs and their putative synapses (with respective colors) are also depicted in **Figure 1**. Overall, our computations suggest that L2/L3-L2/L3 synaptic connection are within a distance of 140 ± 78 μm from the soma.

Next, in order to estimate the magnitude and kinetics of these HL2/L3 - HL2/L3 synapses (see **Methods** **Eqs. (2-5)**), we have used the modeled cell in **Figure 1** to fit the full waveform of the experimental EPSPs (four EPSPs are shown in **Figure 1D**, black traces). The putative synapses at the respective locations for each experimental EPSP was activated, and the synaptic conductance was optimized (see **Methods**) such that the model EPSPs (color traces in **Figure 1D**) would best match the experimental EPSPs. The range of synaptic conductance and kinetics values that fit the HL2/L3 - HL2/L3 excitatory connections is provided in **Table S1**. Assuming five contacts per connection (see **Methods**), the average peak synaptic conductance of HL2/L3 - HL2/L3 connection was 0.88 ± 0.70 nS, with rise time of 0.3 ms and decay time of 1.8 ms. We note that this value characterizes mostly the properties of AMPA-based conductance, as the neuron was recorded at hyperpolarized value (−86 mV) and, thus, for a single connection the NMDA receptors were essentially not activated (see below).

Together, these findings indicate that adjacent human L2/L3 pyramidal cells form synapses with each other predominantly on the proximal basal and oblique dendrites, similar to what was found in the somatosensory cortex of rodents L2/3 pyramidal cells (Sarid et al., 2007, 2013). This finding was further confirmed anatomically in **Figure S1**. Another prediction is that the AMPA-based conductance/contact is rather strong in these cells (0.88 ± 0.70 nS) as compared to 0.3 - 0.5 nS in rodents (Markram et al., 2015; Sarid et al., 2013). Clearly, if more contacts are involved per connection, then this conductance value is overestimated. We will need more detailed reconstructions of connected cell pairs to resolve this uncertainty (See **Discussion**).

### Human dendritic spines with synapses

To obtain a realistic model of a human dendritic spine, we used high-resolution confocal images of spines from HL3 PCs’ dendrites in control post-mortem tissue from human temporal cortex obtained in autopsy (Benavides-Piccione et al., 2013 **Figure 2**, and see **Methods)**. Note that these dendritic spines are much larger than dendritic spines in rodents’ cortical neurons (Benavides-Piccione et al., 2002). From the confocal images, we constructed a prototypical spine model with an average head membrane area of 2.8 μm^2^, spine neck diameter of 0.25 μm (**Figure 2G**), and a spine neck resistance of 50 - 80 MΩ, assuming a specific axial resistance in the spine neck of 200 – 300 Ω-cm. This range of spine neck resistance is in the lower range found in the literature (Svoboda et al., 1996; Tønnesen et al., 2014), see **Figure S2A** and **Discussion.**

The expected effect of excitatory synapses impinging a human dendritic spine was studied by the activation of a spinous synapse, with synaptic properties as found in **Figure 1**, and observing the resultant EPSP in the spine head, spine base and the soma. One such example is shown in **Figure 3B**. We repeated this simulation many times (n = 6,228), by connecting a spine model to each electrical compartment in each of the six modeled human L2/L3 PCs (see cells **in Figure 7** and also in (Eyal et al., 2016)), and then activated a synapse individually at each spine head. The peak EPSP values for all the spines in one of these HL2/L3 models are shown in **Figures 3 C,D**. Activation of a single spinous synapse gave rise to peak EPSP of 12.7 ± 4.6 mV in the spine head membrane, which was attenuated to 9.7 ± 5.0 mV at the spine base and to 0.3 ± 0.1 mV at the soma. Considering each dendritic spine individually, these values represent an attenuation ratio of 1.61 ± 0.93 from the spine head to spine base and 122 ± 196 folds (range of 6 – 1,812 folds) attenuation from the spine head to the soma. This steep attenuations from the spine to the soma is the result of the extended cable structure of human L2/L3 neurons, see (Deitcher et al., 2017; Mohan et al., 2015). Importantly, the small *C_m_* values of ~ 0.5 μm^2^, as we have recently found in human neurons, partially compensated for the otherwise even stronger attenuation. Namely, with *C*_*m*_ of 1 μF/cm^2^, a much steeper voltage attenuation is expected (Figure **3A** in Eyal et al., 2016).

We summarize this section by noting that, locally at the spine head, a depolarization of ~13 mV is expected from an individual excitatory synapse which, on its own, will only minimally activate NMDA-dependent receptors at the spine head. At the soma, a single spiny synapse is expected to give rise to an average peak EPSP of 0.3 mV. The large synaptic conductance in HL2/L3 PCs resulted in a higher peak voltage in the human spine head compared with what we computed for L2/3 rat pyramidal cells (unpublished results). However, due to the strong attenuations in human L2/L3 PCs, the somatic EPSP (from activation of one spine) is similar between the rat and the human (Sarid et al., 2013).

**Figure 3.**
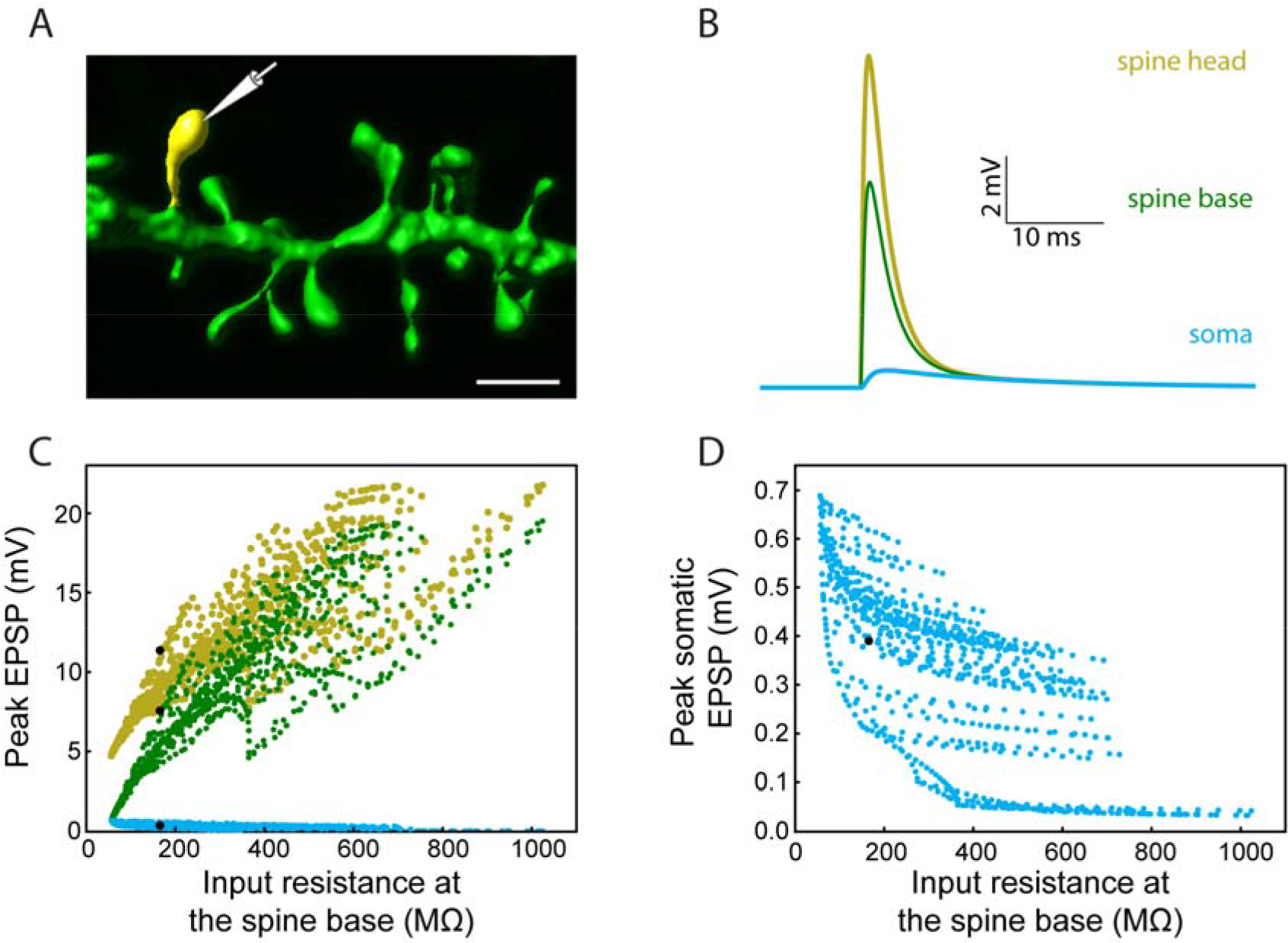
Modeling synaptic inputs on 3D reconstructed human L2/L3 PCs’ dendritic spines. **(A)** A prototypical 3D reconstructed human L3 spine, with average dimensions as found in Figure 2. This dendritic spine was used as a target for excitatory synapses located at its head membrane; synaptic properties are as found in Figure 1. Scale bar = 2 μm. **(B)** Exemplar simulated EPSP in the spine head (yellow), spine base (green) and in the soma (cyan). **(C)** Predicted individual EPSP peak voltage at the (yellow), spine base (green) and in the soma (cyan) for dendritic spines distributed on the modeled cell shown in Figure 1. The spines are arranged according to the input resistance of their respective stem dendrite. Colors are as in **(B)**; black dots depict the example shown in **(B)**. **(D)** Zoom-in into **(C)** showing the peak somatic EPSP.

### Large NMDA currents involved in composite EPSPs; implications for branch-specific NMDA spikes

NMDA-mediated current was shown to be critical for memory consolidation (Shimizu et al., 2000) as well as for computations at the single neuron level, e.g. for shaping the orientation selectivity in the visual cortex (Smith et al., 2013) and for angular tuning in the barrel cortex (Lavzin et al., 2012). These computations were based on the non-linear properties of the NMDA channel. What are the properties of the NMDA channels human neurons? Are human L2/L3 PCs likely to generate local dendritic NMDA spikes similar to those found in rodents?

To experimentally characterize NMDA receptor-mediated currents in HL2/L3 PCs, an extracellular electrode was used for stimulating proximal sites to the apical tree of HL2/L3 PCs while recording the postsynaptic responses from the soma (**Figure 4A**, and **Methods**). The extracellular stimulus gave rise to large composite somatic EPSPs (5.2 ± 1.3 mV, in 3 cells, 6 different stimulation loci). A prominent NMDA-receptor mediated component was isolated after blocking AMPA-, kainite- and GABA_A_-receptors (see also (Wuarin et al., 1992)); three examples from three different HL2/L3 PCs are shown in **Figure 4A**. This data was used to run an exhaustive parameter search, allowing changes in the number of activated synapses and in the dendritic locations of the modeled synapses in search for the AMPA– and NMDA– conductances and kinetics that best fit the experimentally recorded composite EPSP (**Figure 4A**, see **Methods, Eq. (2-6)** for NMDA conductance). The range of parameters for the 100 best models is summarized in **Table S2** and **Figure S3**. The best typical fit among all these acceptable models for this experimental EPSP is depicted in **Figure 4C** (red curves). The corresponding loci of the twenty-one model spinous synapses that provide this fit are shown by the red points superimposed on the dendrite in **Figure 4B**, whereas the parameters for the corresponding AMPA- and NMDA- conductances for this fit are given in **Table S2**. Note the large NMDA conductance amplitude predicted per synapse (1.31 nS) and the large value of γ (γ = 0.077 1/mV compared with 0.062 in (Jahr and Stevens, 1990), see **Eq. (6**)). Large γ values indicate a steep dependence of the NMDA conductance on voltage (**Figure S3B**). Such large NMDA conductance and steepness of the voltage-dependency were also found for most of the other acceptable fits for that experimental EPSP (**Table S2**). Importantly, the value estimated for the AMPA conductance (0.73 nS) is in agreement with the values found in **Figure 1**, although it is based on a different set of data. The same model for AMPA- and NMDA-conductances that was found in **Figure 4C**, was also successful in fitting the other five extracellularly-generated experimental EPSPs (**Figure S4**).

**Figure 4.**
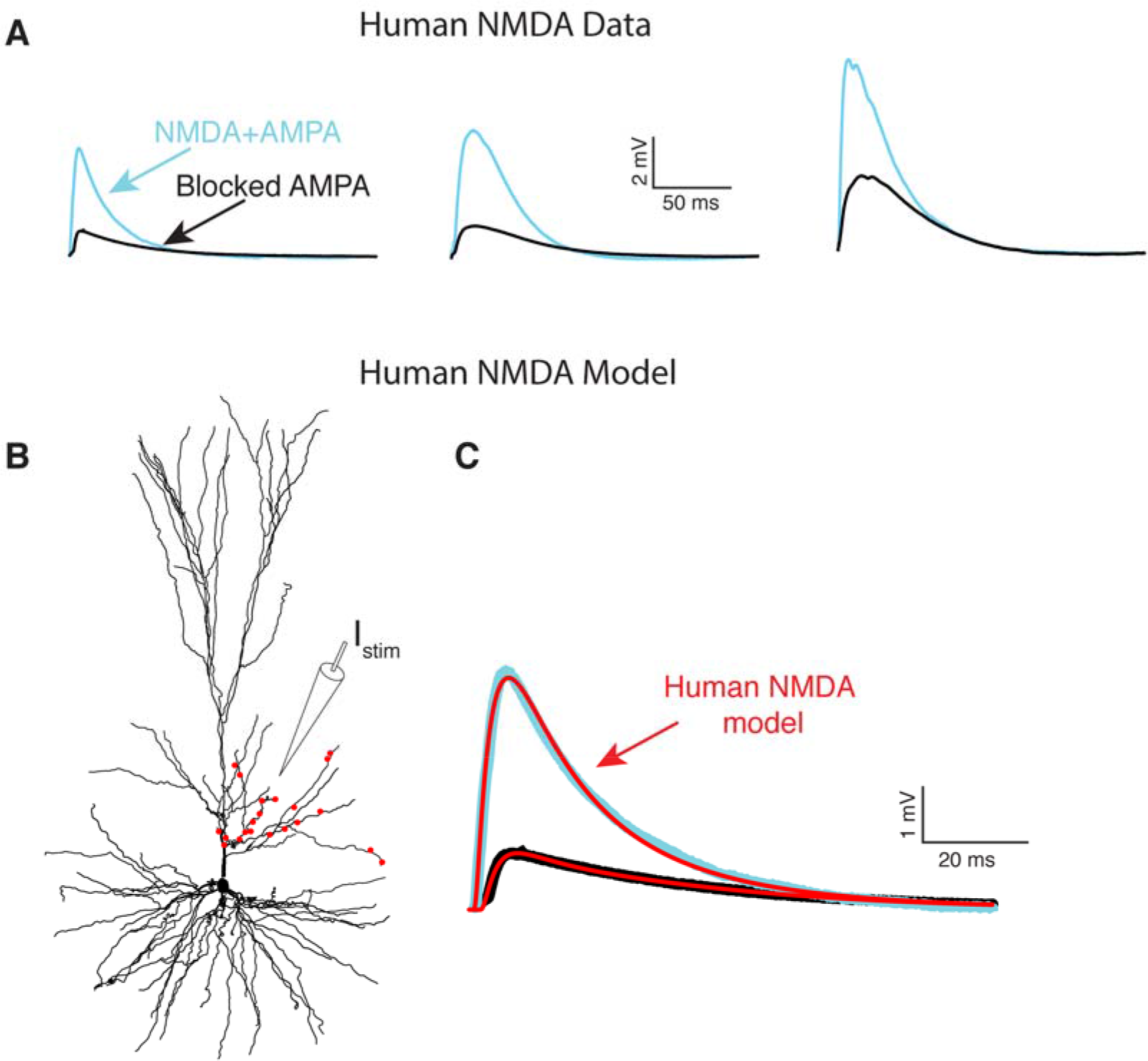
NMDA-receptor based currents in human L2/L3 cells - model fit to experiments. **(A)** Somatic EPSPs recorded in three HL2/L3 PCs in response to stimulation via an extracellular electrode (*I*_*stim*_ in **(B)**). Light blue traces: Somatic EPSPs. Black traces: NMDA-dependent EPSP after blocking AMPA receptors with 1 of NBQX. Leftmost EPSPs were recorded from the cell shown in **(B)**. **(B)** Model prediction for the putative dendritic location and number of activated synapses (red dots) that closely fits the experimental EPSP shown at the top left frame. Synaptic model included both AMPA- and NMDA- based conductances (see **Methods** and **Table S2**). **(C)** Model response at the soma (red traces) when all 21 red synapses in **(B)** were activated. Top trace. EPSP with both AMPA and NMDA conductance; bottom trace, the case in which 82% of the AMPA conductance was blocked (see **Methods**). See similar fits to additional neurons in Figure S4, as well as other accepted models in Figure S3; note the steep non-linearity of the voltage-dependency of the NMDA- current in most of the models (Figure S3C).

What are the implications of such large NMDA-conductance and steep voltage-dependency of the NMDA conductance? We explored this question by inserting for AMPA- and NMDA- based synapses, with the parameters found in **Figure 4C**, into the head membrane of individually modeled dendritic spines as in **Figure 3**. Activation of single axo-spinous synapse with both AMPA- and NMDA- conductances gave rise to EPSP that was generated almost entirely by AMPA current, as the depolarization at the spine head was too small to significantly activate the NMDA-based conductance (see above and **Figure S2B-C**).

Increasing the number of simultaneously activated spiny synapses on a stretch of 20 μm of a basal dendrite (**Figure 5A**) generated an increasingly larger local dendritic EPSP which, for less than 10 synapses, its main source was still the AMPA current (**Figures 5 B,E**). Further increase in the number of activated spines generated supralinear voltage responses due to increasingly stronger recruitment of NMDA-dependent current. Eventually a powerful and prolonged NMDA spike was generated at the dendritic branch (**Figure 5B**), as shown by the corresponding supralinear somatic EPSPs (**Figures 5 D,E**).

Defining an NMDA spike as a local dendritic event with a voltage of at least −40 mV lasting for at least 20 ms, our six model neurons predicted that about 19.9 ± 10.2 simultaneously activated axo-spinous synapses, clustered over 20 μm dendritic stretches, will generate an NMDA spike (n = 5,152). This number decreases to 14.1 ± 7.9 when considering only dendritic terminals (n = 489). These NMDA spikes are spatially restricted (**Figure 5F**), much due to the unique properties of human basal dendrites which are both highly branched and, importantly, terminate with a particularly electrically elongated branch (unparalleled in L2/3 pyramidal cells of mouse and rats (Deitcher et al., 2017; Mohan et al., 2015)). The emergence of local NMDA dendritic spikes was demonstrated in several experimental and in theoretical studies in rodent pyramidal neurons (Major et al., 2013; Palmer et al., 2014; Schiller et al., 2000; Schmidt-Hieber et al., 2017; Smith et al., 2013). Interestingly, although human L2/L3 pyramidal neuron dendrites, especially the basal terminals, are electrotonically more extended than those in rodents (Deitcher et al., 2017; Mohan et al., 2015), a similar number of NMDA-based synapses were required to generate a local NMDA spike. This results from a combination of the relatively large NMDA-conductance in HL2/L3, its steep voltage-dependency, and the low *C*_*m*_ value in these cells which promotes local excitability/nonlinearity (Eyal et al., 2016), see **Discussion**.

### Multiple NMDA-based nonlinear functional subunits in human L2/L3 pyramidal cells

The larger the number of nonlinear dendritic subunits, the larger the storage capacity of a neuron (Poirazi and Mel, 2001). Given the spatial restriction of the NMDA spikes in HL2/L3 pyramidal cells and the large number of dendritic branches/terminals in these cells, we seek to quantify the potential number of independent NMDA-spikes (functional dendritic subunit) in these cells. The definition for an independent NMDA spikes/independent functional dendritic subunit is provided in the **Methods** section.

We started by using Rall’s cable theory to analyze the degree to which distal basal trees in HL2/L3 PCs are electrically decoupled from each other. In electrically decoupled dendrites, local nonlinear events are likely to be independent (affect each other minimally). **Figure 6A** shows the remarkable electrical isolation of the distal basal dendrites in these cells (blue zone), as captured by *R*_*i,j*_ the transfer resistance from branch *i* to branch *j*. The smaller *R*_*i,j*_ is, the larger the voltage attenuation from *i* to *j* and the larger the electrical decoupling between these two locations. The functional consequence of this significant electrical decoupling between distal basal dendrites is manifested in the capability of these dendrites to generate many independent NMDA spikes simultaneously (**Figures 6 B,C**). A comprehensive search for the space of possible independent simultaneously activated NMDA spikes in HL2/L3 PCs models provided an estimate of 24.8 ± 4.4 (n = 6) independent NMDA spikes per cell (**Figure S5**) as compared with only 13.7 ± 2.1 (n = 3) in rat L2/3 pyramidal cell model (not shown, models are based on (Markram et al., 2015; Sarid et al., 2013)). This increased number of local nonlinear dendritic subunits in HL2/L3 PCs enhances the memory/computational capacity and computational capabilities of these cells (see **Discussion**).

### Active axo-somatic models of human L2/L3 PCs

Next, we focused on constructing models for the somatic spiking properties of HL2/L3 PCs. Towards this end, repeated supra-threshold depolarizing current steps were recorded from the six HL2/L3 PCs described above (example traces are shown in **Figure 7A**, black traces). This data was complemented with I-F curves recorded in 25 other human L2/L3 PCs (grey traces in **Figure 7B**; the average I-F curve for human L2/L3 PC is shown in black). From these experimental spikes, we extracted a set of characteristic features (spike width, height, frequency, adaptation index etc., see **Table S3**), and employed multiple objective optimization (MOO) to fit these experimental features via conductance-based neuron models (Druckmann et al., 2007; Hay et al., 2011). This procedure yielded a good fit between models and experiments (**Figure 7**, color traces). The values for the membrane ion channels involved in generating these six HL2/L3 models are provided in **Table S4.** These modeled cells are available for download in modelDB (http://modeldb.yale.edu/238347).

### Relatively small number of L2/L3 - L2/L3 synapses ignite somatic Na^+^ spike

Now that we have a faithful model for the spiking activity (and for the spike threshold) in HL2/L3 PCs, we may ask how many excitatory axo-spinous synapses should be simultaneously activated for initiating a somatic Na^+^ spike in these cells? Two cases were tested: in the first case, the excitatory (AMPA- and NMDA- based) synapses were randomly distributed over the modeled HL2/L3 dendrites (**Figure 8A**, green curve for one of the modeled cells and green column in **Figure 8B** for the six modeled cells). In the second case the synapses were clustered such that local dendritic NMDA spikes were likely to occur (**Figure 8A**, pink curve and pink column in **Figure 8B**).

In both cases, a similar number of excitatory synapses were required for generating a somatic spike with 50% probability, 133 ± 16 (n=6) synapses in the distributed case and 136 ± 38 in the clustered case. However, the clustered case is shallower, implying that in some instances fewer synapses were sufficient for generating a somatic spike (left part of the pink curve in **Figure 8A**) and in some cases more synapses where required to generate a somatic spike (right part of the pink curve in **Figure 8A**, see also Farinella et al., 2014). In the former cases, the clustered inputs (with their corresponding prominent NMDA current) were located at proximal branches and in the latter case they were clustered either at distal branched (as in **Figures 6 B,C** and see also examples in **Figure S6A)** or in branches with small input resistance, such that the threshold for NMDA spike was not reached. In other cases, two or more clusters were located on the same branch leading to saturation of the local membrane voltage (**Figure S6A**). Specific examples for suprathreshold clustered versus distributed inputs are shown in **Figure 8C**. Note that clustered input has a larger probability of generating a burst of somatic spikes due to the prolonged duration of the dendritic NMDA spikes.

It is interesting to note that the number of excitatory synapses that were necessary for generating a somatic spike in human L2/L3 PCs is not significantly larger than what we predicted for the electrically more compact L2/3 PCs of the rat; ~125 synapses for the distributed case and ~145 synapses for the clustered case in a modeled cell from (Markram et al., 2015); see **Figure S6B**. This is explained by the more potent excitatory synapses (larger AMPA and NMDA-based conductances) in human L2/L3 pyramidal neurons compared to rodents (Sarid et al., 2013) and the smaller *C*_m_ value in human L2/L3 PCs (Eyal et al 2016). Therefore, a relatively small number of simultaneously activated excitatory synapses are sufficient for reaching threshold for spike firing at the soma/axon of human L2/L3 PCs. Importantly, the estimated number of dendritic spines/neuron, when considering only the basal trees of HL2/L3 pyramidal cells in the prefrontal and temporal human cortices, is 15,138 and 12,700, respectively (Elston et al., 2001); these numbers agree with measurements of spine density (Benavides-Piccione et al., 2013) and with our measurements of total dendritic length of 3D reconstructed human L2/L3 PCs (Mohan et al., 2015). Based on the above, we estimate that the total number of dendritic spines in HL2/L3 PCs ranges between 20,000 - 30,000. With only about 135 simultaneously activated excitatory synapses required to fire an axonal spike, the number of combinatorial synaptic possibilities for generating axonal output in HL2/L3 PCs is enormous.

We summarize the above two sections by highlighting that our first-ever realistic model of the somatic spiking activity in human cortical L2/L3 neurons, together with the biologically-based model estimate of the strength of excitatory (AMPA + NMDA − based) synapses in these cells, enabled us to demonstrate that (i) around 20 clustered spinous synapses are likely to generate a local dendritic NMDA spike and (ii) that a somatic Na^+^ spike is likely to be generated at the soma/axon region when a rather small number of these synapses are activated simultaneously. These findings are surprising as human L2/L3 neurons are much larger (in both the total dendritic length/surface area and in the number of dendritic branches) than the respective cells in rodents. Still, due to various compensatory mechanism (small *C_m_* value and large *R*_*m*_ value (Eyal et al., 2016) and more potent excitatory synapses), similar numbers of excitatory synapses ignite both the local NMDA spike and the somatic spike in L2/L3 pyramidal neurons of humans and rodents. This issue will be further elaborated in the **Discussion**.

## Discussion

The structure and function of local neuronal circuits, with their specific cell types and highly selective connectivity pattern, are presently in the focus of worldwide efforts (Egger et al., 2014; Hawrylycz et al., 2016; Kasthuri et al., 2015; Markram et al., 2015). In the midst of this effort star single neurons, the elementary processing units in the brain, that, due to their complex morphological and membrane nonlinearities, function as sophisticated computational and plastic units (Rall, 1959; review in Stuart et al., 2016). Of particular interest are cortical pyramidal cells; these principal neurons represent the majority of neurons composing the mammalian cortex, they are the major source of excitatory cortical synapses and their dendritic spines are the main postsynaptic target of excitatory synapses. Pyramidal cell dendrites exhibit highly nonlinear properties, including local NMDA spikes in distal dendritic branches and more global Ca^2+^ spikes in the apical tuft. These dendritic nonlinearities impact both local plastic processes and shape the spiking output at the axon and, consequently the dynamics of cortical networks (Bono et al., 2017; Hay and Segev, 2014; Larkum et al., 1999, 2009; Major et al., 2013; Mel et al., 2017; Poirazi et al., 2003a; Smith et al., 2013; Spruston, 2008). All these studies, focusing on the computational consequences of dendritic complexities, were performed on rodents (mouse and rat). What could be learned from those on human neurons remained an open question, because we lacked similar systematic studies on cortical pyramidal cells (or on any other neuron type) in the human neocortex.

The present study attempted to narrow this gap by linking experimental data from human cortical pyramidal cells with detailed models of these same cells. Towards this end we integrated in our detailed models a variety of properties of L2/L3 pyramidal cells from human temporal cortex, including dendritic morphology and physiology, dendritic spine anatomy, synaptic properties and somatic spiking characteristics. Our overarching approach provided several new insights on integrative properties of HL2/L3 PCs and on their computational capability.

### Properties of human L2/L3-L2/L3 excitatory synapses

Our HL2/L3 PCs models predicted larger conductance of both AMPA- and NMDA- receptors as compared to rodents. This is expected as the spine head surface area is larger in human neurons (Benavides-Piccione et al., 2002; DeFelipe et al., 2002). In human L2/L3 PCs, the average estimated AMPA conductance is ~0.8 nS per contact (**Figures 1** and **4** and **Tables S1,2**) as compared to 0.3 - 0.5 nS in rodents; the NMDA conductance is estimated to be ~1.3 nS per contact (**Figure 4** and **Table S2**) as compared to ~0.4 nS in rodents (Markram et al., 2015; Sarid et al., 2007, 2013). We note that all the above estimates are based on the assumption that single axon connecting HL2/L3 neurons to each other make, on average, 5 synaptic contacts, similar to the case in rodents (Feldmeyer et al., 2002, 2006; Markram et al., 2015). This estimate requires further study. Still, the total conductance for a HL2/L3-HL2/L3 PCs connection is about twice as strong, on average, than in the respective rodents’ synapse.
We used cable theory and Rall’s “shape indices” (Rall 1967) to estimate the electrotonic locus of HL2/L3-HL2/L3 PCs synapses. The fast rise-time and the relatively narrow half-width of the respective somatic EPSP indicated that these connections are made at the proximal dendrites (**Figure 1**), as indeed was further validated by the reconstruction of L2/L3-L2/L3 putative synapses (**Figure S1**). Similar findings hold for rodents as well, whereby connected pairs of L2/3 PCs (as well as of L5 PCs), make proximal synapses with each other (Markram et al., 1997a; Sarid et al., 2013). We note that our model predicts that the NMDA current is essentially not activated by a single HL2/L3- HL2/L3 synapse (**Figure 5**).

### Dendritic NMDA spikes in human PCs

NMDA conductance is voltage-dependent; the steepness of this dependency is determined mainly by the parameter γ in **Eq. (6**). Fitting the experimental data to the model provided an estimate of γ ~0.075 (**Figure 4** and **Figure S3**), which is larger (steeper voltage dependency) than in rodents pyramidal cells (Jahr and Stevens, 1990), implying that human L2/L3 PC dendrites are prone to generating a local dendritic NMDA spike. Indeed, our model shows that, on average, about 20 ± 10 simultaneously activated excitatory spinous synapses, stretched over 20 μm of dendritic branch, are likely to ignite large and prolonged NMDA spikes (**Figure 5**). At distal dendritic terminal branches, due to their large input resistance (up to 1GΩ), only 14 ± 8 synapses were required for generating an NMDA spike. Similar numbers of excitatory synapses were required for generating branch-specific NMDA spikes in models of rodents’ dendrites (Farinella et al., 2014; Larkum et al., 2009; Rhodes, 2006). We conclude that human L2/L3 PC’s dendrites are prone to generating local NMDA spikes, both due to the properties of the NMDA-conductance as well as due to the low *C*_*m*_ and large *R*_*m*_ values in these cells (Eyal et al., 2016).

**Figure 5.**
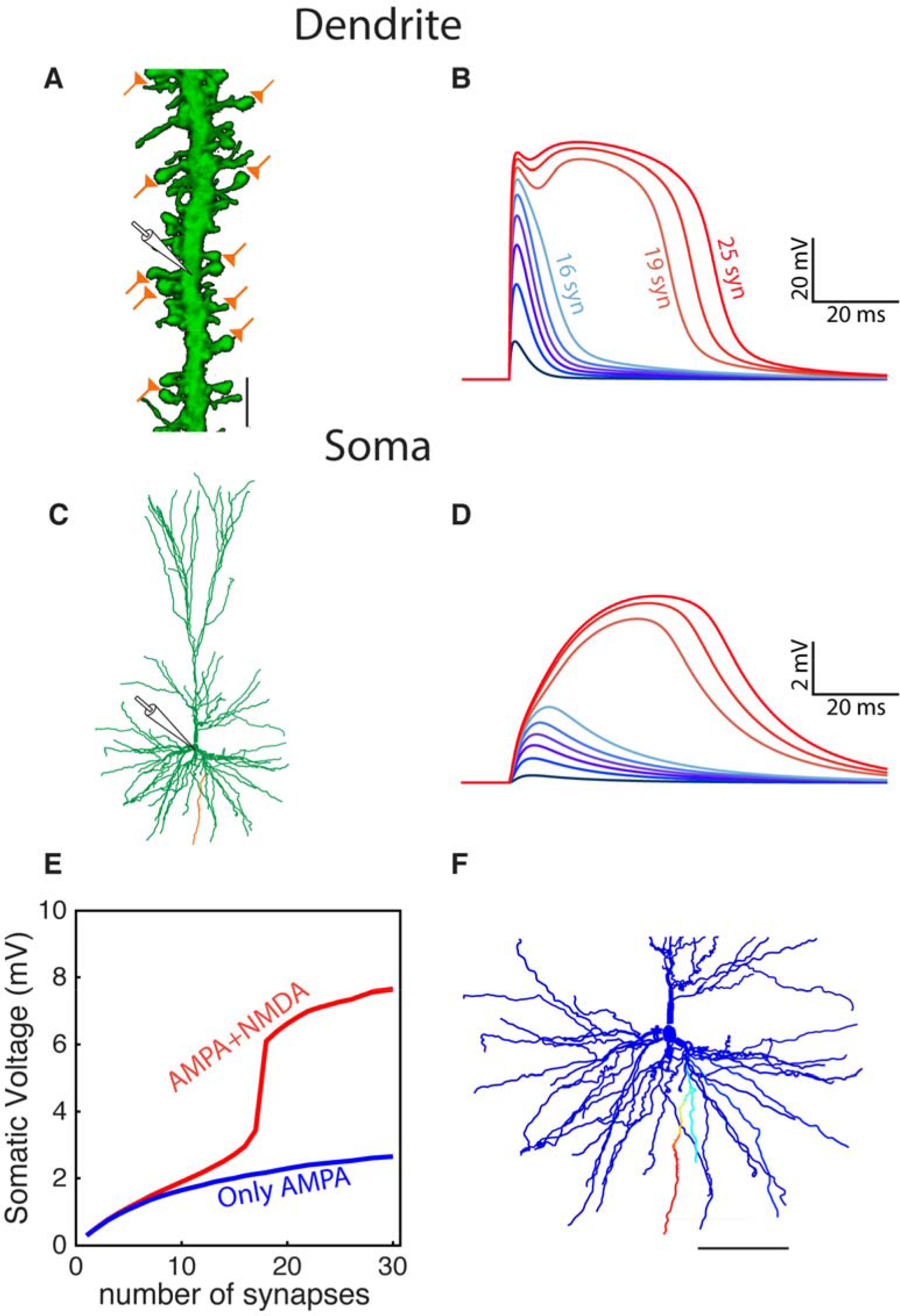
Modeled dendritic NMDA spike in distal dendrites of HL2/L3 neurons. **(A)** Confocal image of a dendrite from human L2/L3 pyramidal neuron (obtained from postmortem preparation, see **Methods**) that is densely decorated with dendritic spines. The location of the activated model synapses is illustrated by the “orange synapses” (scale bar = 5 μm) that were simulated on a similar basal dendrite from the modeled HL2/L3 cell in **(C)** (orange branch). **(B)** Voltage response at the stem dendrite when increasing the number of simultaneously activated spine synapses. Activated synapses are distributed within 20 μm of dendritic stretch. Note the steep nonlinear change in local dendritic voltage when 19 synapses were activated - resulting in an NMDA spike. **(C)** The morphology of the modeled cell. **(D)** Somatic voltage in response to synaptic activation as in **(B). (E)** The somatic EPSP amplitude as a function of the number of activated dendritic synapses with NMDA (red) and when only AMPA current was activated (blue). **(F)** The spatial extent of the NMDA spike in one basal dendrite - voltage is color-coded (blue, -86mV, red, -10 mV; scale bar = 100 μm). The NMDA spike was activated by 20 clustered synapses and the voltage was recorded 10 ms after their synchronous activation. Note the large number (44) and the distinctive elongation of the basal terminals in this cell.

### Models of human dendritic spines

This is the first study that models human dendritic spines. Towards this end, we employed a large high-resolution confocal images and 3D reconstructions of dendritic spines from HL2/L3 PCs (**Figure 2**). From this data set we constructed a prototypical human spine model to explore the range of EPSPs values of spinous synapses (**Figures 2, 3**) and to estimate the spine neck resistance. Activation of single spinous generated EPSP with peak voltage of 12.7 ± 4.6 mV in the spine head membrane, 9.7 ± 5.0 mV at the spine base and 0.3 ± 0.1 mV at the soma. These values represent a mean attenuation ratio of ~1.6 from the spine head to spine base and ~120 folds attenuation from the spine head to the soma, which results from the extended cable structure of human L2/L3 neurons (Deitcher et al., 2017; Mohan et al., 2015). The small *C*_*m*_ values of ~ 0.5 μm^2^ in human neurons and the corresponding large *R*_m_ value in these cells (Eyal et al., 2016) partially compensated for the otherwise even steeper dendritic attenuation.

The above value of the single spinous EPSP relies on our estimates of the spine neck resistance which, due to the relatively thick neck in human spines (~ 0.25 μm), was estimated to range between 50 – 80 MΩ. This estimate is based on the assumption that the specific axial resistivity of the spine neck is 200-300 Ωcm, as was the value for the dendrites in our models (see discussion in Eyal et al., 2016). It could be the case that, due to various intracellular filaments/organelles in the spine neck, the effective axial resistance is larger than assumed here. Small variations in the spine neck diameter (e.g., ± 0.05 μm) or in the axial resistivity (e.g., ± 100 Ω-cm) would lead to spine neck resistance ranging between 19 – 128 MΩ (**Figure S2A**).

Recent studies used a variety of methods to directly estimate the spine neck resistance in rodents. These estimates vary significantly between different groups, from around 50 MΩ (Popovic et al., 2015; Svoboda et al., 1996; Tønnesen et al., 2014) to 0.5-3GΩ (Araya et al., 2014; Harnett et al., 2012; Palmer and Stuart, 2009). Note a recent study by (Kwon et al., 2017), with estimates of 101 ± 95 MΩ for the spine neck resistance in cultured hippocampal neurons from mice, using a voltage indicator and glutamate uncaging.

### Models for somatic/axonal Na^+^ spikes in human L2/L3 PCs

We used a large dataset of ~900 spikes recorded from six HL2/L3 PCs to model the axonal/somatic Na spiking activity in these cells (**Figure 7)**. Towards this end we used our multiple objective optimization method (Druckmann et al., 2007; Hay et al., 2011) to estimate the conductance of a set of membrane ion channels, assuming that the kinetics of these ion channels are similar to that of rodents (Ranjan et al., 2011). Our models predicted that the *Na*^+^ - channels density in the axon of HL2/L3 PCs is relatively high (4.9 ± 1.0 S/cm^2^, n=6, see review in (Kole and Stuart, 2012)). This value results from the need to fit the large peak of the Na^+^ spikes in these neurons (38.6 ± 3.2 mV, from a voltage base of −83.5 ± 2.9 mV) and to compensate for the large current sink imposed on the axons’ initial segment by the (huge) dendritic tree in human L2/L3 neurons. Similar densities were estimated recently also for L2/3 and L5 PCs in the rat (Markram et al., 2015). The high densities of the sodium channels go along with large densities of Kv3.1 ion channel (1.9 ± 0.1 S/cm^2^), resetting the strong inward sodium currents and yielding the narrow spikes measured in these cells (half width of 1.02 ± 0.15 ms). On the other hand, the densities of the persistent potassium were low both for the axon (0.019 ± 0.039 S/cm^2^) and for the soma (0.0002 ± 0.0004 S/cm^2^), allowing the models to reach high firing rates of 20-30 Hz for strong current inputs (**Figure 7B**). High densities for sodium channels were required for the somatic compartment as well (0.33 ± 0.17 S/cm^2^). Note also that, in rodent PCs, the dendritic membrane is endowed with low density of sodium channels, effectively decreasing the current sink due to the dendrites and supporting the back-propagating action potential (Stuart et al., 2016; Stuart and Sakmann, 1994). We still lack recordings from human PCs’ dendrites and, consequently, do not know if and to what extent HL2L/3 PC’s support back-propagating APs. Yet, at least for the spiking activity recorded at the soma, our models seem rather faithful (**Figure 7**).

### From spiny synapses to dendritic and axonal spikes

Our comprehensive HL2/L3 PCs models, integrating both dendritic morphology, synaptic properties, spine properties and somatic active ion channels, provided an estimate for the number of simultaneously activated HL2/L3 – HL2/L3 synapses required to generate a somatic Na^+^ spike (**Figure 8**). We found that about 135 synapses are required to fire a somatic/axonal spike with a probability of 50%, both for a spatially distributed case as well as for a case in which the excitatory synapses were clustered (**Figure 8**). We note that, although similar number of synapses are required to generate a somatic/axonal spike in both cases, the span of the curves is significantly different. In the distributed case, the probability for generating a spike is a steep function of the number of synapses. Indeed, for the model shown in Figure **8A,** the probability to generate a spike with less than 110 synapses is very low whereas with 140 a spike will always be generated. In contrast, the clustered case is more elongated and it has a larger variance. For example, in some cases as few as 80 synapses (four clusters with 20 synapses each) were sufficient to generate spike, and in other cases as much as 200 synapses were not enough (**Figure 8**, **Figure S6**). These clustered inputs enable the cell to have a larger dynamic range for both I/O relationship as well as for NMDA-dependent calcium-based plasticity, see discussion below.

Interestingly, the number of excitatory synapses required to generate a somatic spike was similar for the HL2/L3 PCs models as for the rodent L2/3 PC model (**Figure S6**), even though the human L2/L3 PCs are much larger (both anatomically and electrotonically, (Mohan et al., 2015). Similarly, about the same number of clustered dendritic synapses are required to generate an NMDA spike in both human and rodent neurons. This is explained by two compensatory mechanisms in HL2/L3. The reduced *C*_*m*_ value and increased *R*_*m*_ value (Eyal et al., 2016) enables the cells to charge the membrane more effectively. Furthermore, the large AMPA- and NMDA- conductances in these cells also compensates for the increased size of HL2/L3 cells. Therefore, the synapses-to-spike ration is preserved in both rodents and human L2/L3 PCs. The somatic spike “requires” around 135 excitatory synapses and the local NMDA spike about 20 excitatory synapses.

Yet, due to the large number of HL2/L3 terminal basal branches (44.5 ± 8.1, n=6) and their particular cable elongation, HL2/L3 are functionally distinctive as compared to rodents L2/3 PCs (where the number of basal terminals is 31.4 ± 8.6, n=15, see (Deitcher et al., 2017; Mohan et al., 2015). These two distinctive features in HL2/L3 PCs result in significant electrically decoupling of the basal terminals from each other (**Figure 6),** enabling HL2/L3 PCs dendrites to function as multiple, semi-independent (nonlinear), subunits. Indeed, our models predicted that HL2/L3 distal could generate ~25 simultaneous and independent local NMDA spikes, particularly at the basal dendrites, versus a total of 14 independent NMDA spikes in rat PCs.

**Figure 6.**
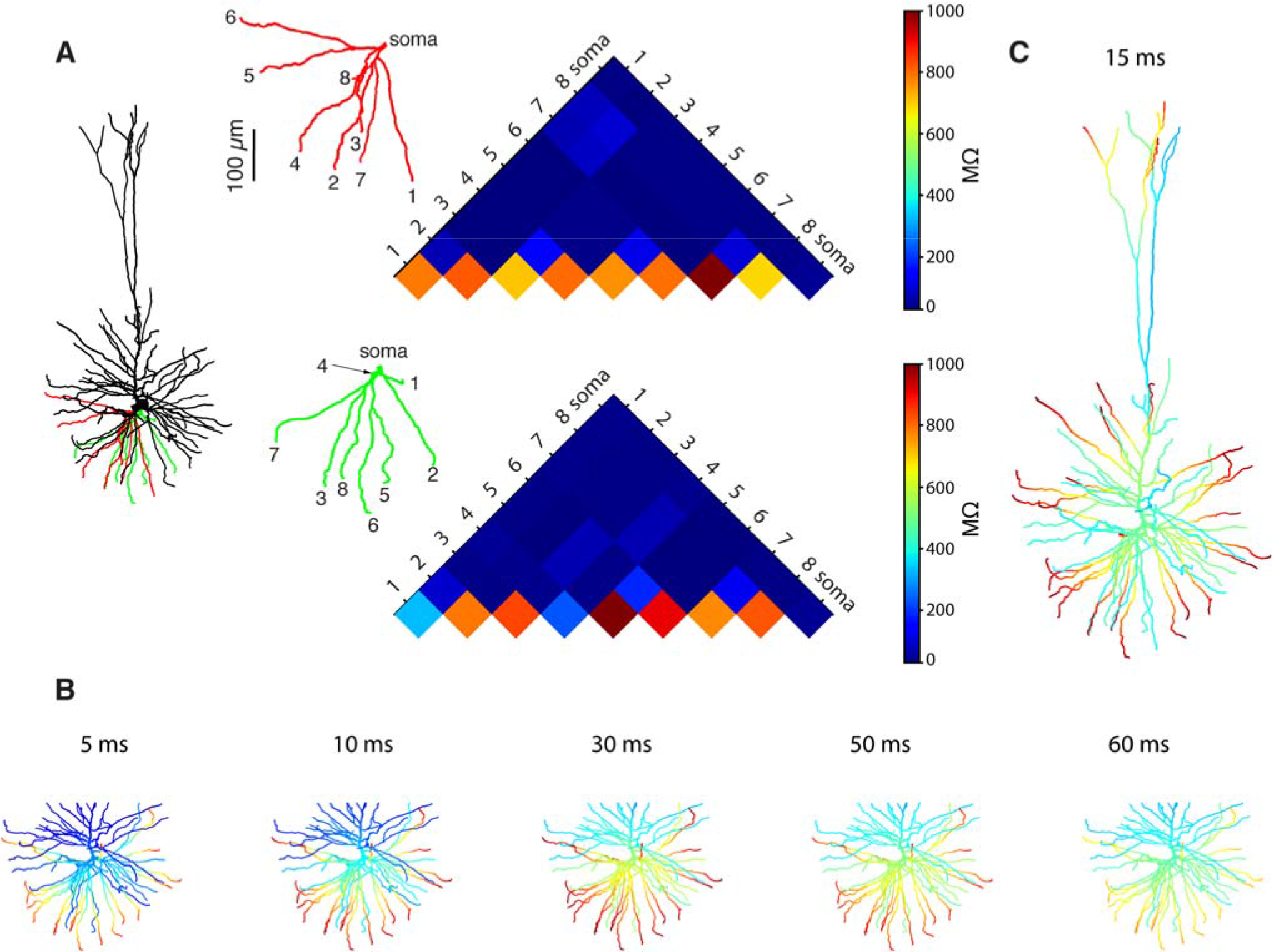
Multiple NMDA-based functional subunits in human L2/L3 pyramidal cells. **(A)** Significant electrical decoupling of the basal dendrites from each other. Left, red and green depict two basal subtrees taken from the modeled cell shown at left. Right, color-coded matrix showing the transfer resistance (*R*_*i,j*_) between one tip of a basal terminal *i* and the tip of the other basal terminal *j*, for the red and green basal subtrees. Blue colors represent small transfer resistance (significant electrical decoupling); the lower part of the triangles (hot colors) depicts the input resistance (Rμ) at the different dendritic tips. **(B)** Twenty-one independent simultaneous NMDA spikes could be generated in the modeled basal tree (red branches, see **Methods**). Clusters of excitatory synapses that were sufficient to generate local NMDA spikes were activated simultaneously at t = 0 ms and the membrane voltage (color coded) as a function of time is superimposed on the simulated dendritic tree. **(C)** When activating the entire dendritic tree with clustered synaptic inputs, twenty-eight independent NMDA spikes could be generated simultaneously in the modeled L2/L3 neuron (red dendritic terminal branches, basal plus apical trees). See **Methods** for the definition of “an independent NMDA spike”.

### Assumptions and missing data

It is important to note that despite the amount of growing research, there are still a small number of biophysical and anatomical studies on human neurons cells (see however recent release by the Allen Institute, http://celltypes.brain-map.org/). Consequently, our study is based on several assumptions. Particularly, our estimates for the synaptic parameters (**Tables S1, S2**) are based on the assumption that HL2/L3 - HL2/L3 PCs connections are formed by five synaptic contacts. Multiple contacts per connection is typical for cortical excitatory neuronal synapses (e.g., (Markram et al., 1997b; Sarid et al., 2007, 2013)) and is supported by **Figure S1** also for human cells. Interestingly a recent paper (Molnár et al., 2016) showed that the number of light microscopy detected synaptic contacts between human pyramidal neurons and human inhibitory neurons is ~3 per connection, similar to what was found in rats. However, in human neurons, each one of the presynaptic active zones contained about 6 functional release sites comparing with only about 1.5 in rats. This agrees with our prediction of stronger synapses per contact in human excitatory synaptic contact. Yet, a study similar to (Molnár et al., 2016) for human PCs-PCs is required for assessing the number of synaptic contacts per connection.

In addition, the estimates in **Figure 8** for the number of synapses required to generate a dendritic NMDA spike and a somatic *Na*^+^ spike is based on the assumption that all the excitatory inputs to the HL2/L3 PCs have similar properties to those activated by an input from a proximal HL2/L3 neuron as in **Figure 1**, and to those activated by an extra cellular electrode located near the apical tree as in **Figure 4**. Variations in the synaptic properties as function of the location of the synapse on the dendritic tree (Magee and Cook, 2000) and as function of the source of the input (Sarid et al., 2007, 2013) were found in mammals and are probably expected also in the human brain. A more complete database, studying the synaptic properties of different connections in the human neocortex is therefore required in order to improve the validity of our predictions.

**Figure 7.**
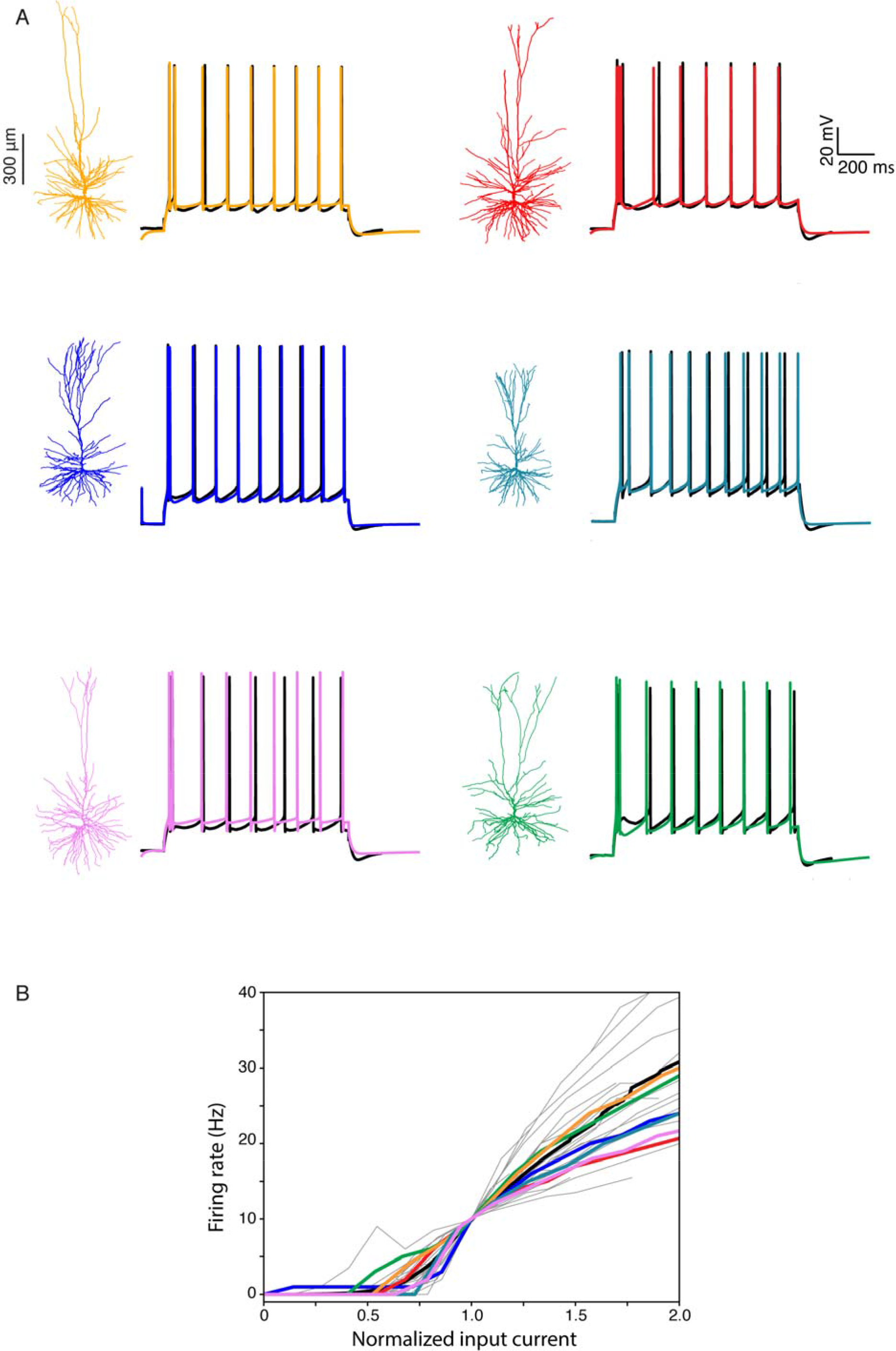
Modeling somatic/axonal Na^+^ spikes for six human L2/L3 PCs. **(A)** Fit between models and experiments. Black traces, experimental spike trains recorded from human L2/L3 PCs shown on the left. Step current input was selected to generate spike train of about 10 Hz. Color traces, model responses to the same experimental current step. Models were optimized using multiple objective optimization algorithm (MOO, see **Methods**). **(B)** Grey traces, experimental I-F curves in 25 human L2/L3 PCs, normalized by the input current corresponding to a firing rate of 10 Hz. Black curve, average of all experimental traces. Colored traces, theoretical I-F curves for the six modeled cells shown in **(A)** with corresponding colors.

Another missing experimental component is the properties of ion channels at the soma/axon region in human neurons. We have successfully used models of ion channels from rodents (see recent intense effort to build such models for rodents’ cortical neurons based on ion channel characterization – the Channelpedia (http://channelpedia.epfl.ch/) (Ranjan et al., 2011)). Yet, we are aware of the fact that membrane ion channels may differ between species (Angelino and Brenner, 2007), although a comprehensive study by Ranjan et al., (in preparation) highlights the great similarity between human and rodents ion channel. Finally, as mentioned above, dendritic recordings from human dendrites are still missing; this will be essential for validating our prediction that HL2/L3 PCs are prone to generating multiple branch-specific NMDA spikes.

### Functional significance of the morpho-electrical complexity of human L2/L3 dendrites

Following the above results, we hereby propose a new index for defining the functional complexity of cortical neurons based on the maximal number of independent-simultaneous NMDA spikes that the neuron could generate (see also, Koch et al., 1982; Mel, 1992; Poirazi and Mel, 2001; Polsky et al., 2004). Note that, for this definition, we do not assume *a priori* that different dendritic branches are functionally independent as in (Poirazi and Mel, 2001, see discussion in Jadi et al., 2014; Poirazi et al., 2003b;). Rather, the number of subunits is determined by the degree (or lack thereof) of interaction between the dendritic branches. This is determined by the number of the branches and the cable structure of the tree (**Figure 6A**). It is important to note that our definition of the cell’s complexity index provides a lower bound, as if the NMDA spikes were not activated simultaneously, then the number of the nonlinear dendritic subunits would be much larger, in the order of the total number of branches per cell. Indeed, our definition takes the strictest constrain. Namely, that the NMDA-spike should not interact with each other when activated simultaneously. Clearly, the estimated number of independent NMDA spikes is also determined by the “state” of the neuron; whether more shunted via being bombarded by background active synapses as in the *in vivo* case or more quiescent as in the *in vitro* case as we used (Farinella et al., 2014).

Using the definition described above, we found that HL2/L3 PCs have ~25 nonlinear dendritic subunits, compared with only ~14 in the rat. This suggests that HL2/L3 PCs have augmented computational capabilities compared with the rodent L2/3 PCs. (Poirazi and Mel, 2001) provided a measure for the storage capacity of neurons based on their structural and nonlinear dendritic properties. Upper bounds on the capacity were derived for two cases: the case of a linear neuron, where the cell activity is determined by a linear weighted sum of all its inputs, similar to a one-layer neural network (McCulloch and Pitts, 1943), and the case in which nonlinear synaptic integration was first implemented in a given dendritic subunit and then summed-up to produce the axonal output (2 layer model). They then computed the storage capacity of the neuron for these two cases, see eq. (4) in (Poirazi and Mel, 2001). Using our definition for the number of independent nonlinear dendritic subunit and taking into account that human L2/L3 PCs have about 3-times more synapses/cell (Ballesteros-Yáñez et al., 2006; DeFelipe et al., 2002; Elston et al., 2001) we computed the storage capacity of HL2/L3 PCs to be 23.4*10^4^ bits compared with 6.3*10^4^ bits in respective cells of the rat (**Figure 9**).

The analysis described above is based on the structural plasticity of synapses. Namely on the number of ways to connect presynaptic axons with the post synaptic neuron. However, the memory capacity of a neuron can also be studied for a given connectivity. The neuron may learn to respond (or not respond) to specific patterns by assigning different weights to its synapses (Mel et al., 2017). The number of patterns that could be learned by a neural network with E parameters (edges in the network) is O(E*log(E)) for networks with sign activated neurons and O(E^2^) for networks with sigmoid output neurons (Shalev-Shwartz and Ben-David, 2014). Consequently, having both more synaptic inputs in HL2/L3 PCs (a larger input layer) and more nonlinear dendritic subunits (larger second layer) increases E and thus dramatically increases the computational capacity of these neurons.

Clearly, the input-output properties of neurons could only be approximated by one- or two- layer networks (Poirazi et al., 2003b). This is apparent by observing the large variance in the number of clusters required to generate AP in **Figure 8**. When clustered inputs are impinging in proximal dendrites, the depolarization will effectively spread to other subunits (Jadi et al., 2014). Furthermore, the present study assumed the extreme cases where inputs were activated simultaneously and were either clustered or uniformly distributed over the dendritic surface (**Figure 8)**. Obviously, clustered inputs interact with distributed background activity and the temporal variance in the input has a large impact on the neuron’s output (Farinella et al., 2014). To replicate the I/O properties of the neuron under such realistic conditions, a more complex perhaps “deep network” might be required. This remains an interesting challenge for future studies.

In summary, the present work is the most comprehensive, experimentally-based, modeling study of any human neuron. To fully model human neurons, more data than we presently have is required (e.g., the properties of inhibitory and modulatory inputs and of additional dendritic excitably, such as the back-propagating AP and the dendritic Ca-spike). Still, the present study significantly increases our acquaintance with human pyramidal cells, laying the foundation for constructing models for other (excitatory and inhibitory) neuron types in the human brain paving the way for constructing models of human cortical microcircuit and exploring its dynamical repertoire *in silico,* as has been recently performed for rodents cortical microcircuits (Egger et al., 2014; Markram et al., 2015). This will bring us closer to understanding “what makes us human” – at least at the scale of cortical microcircuits.

**Figure 8.**
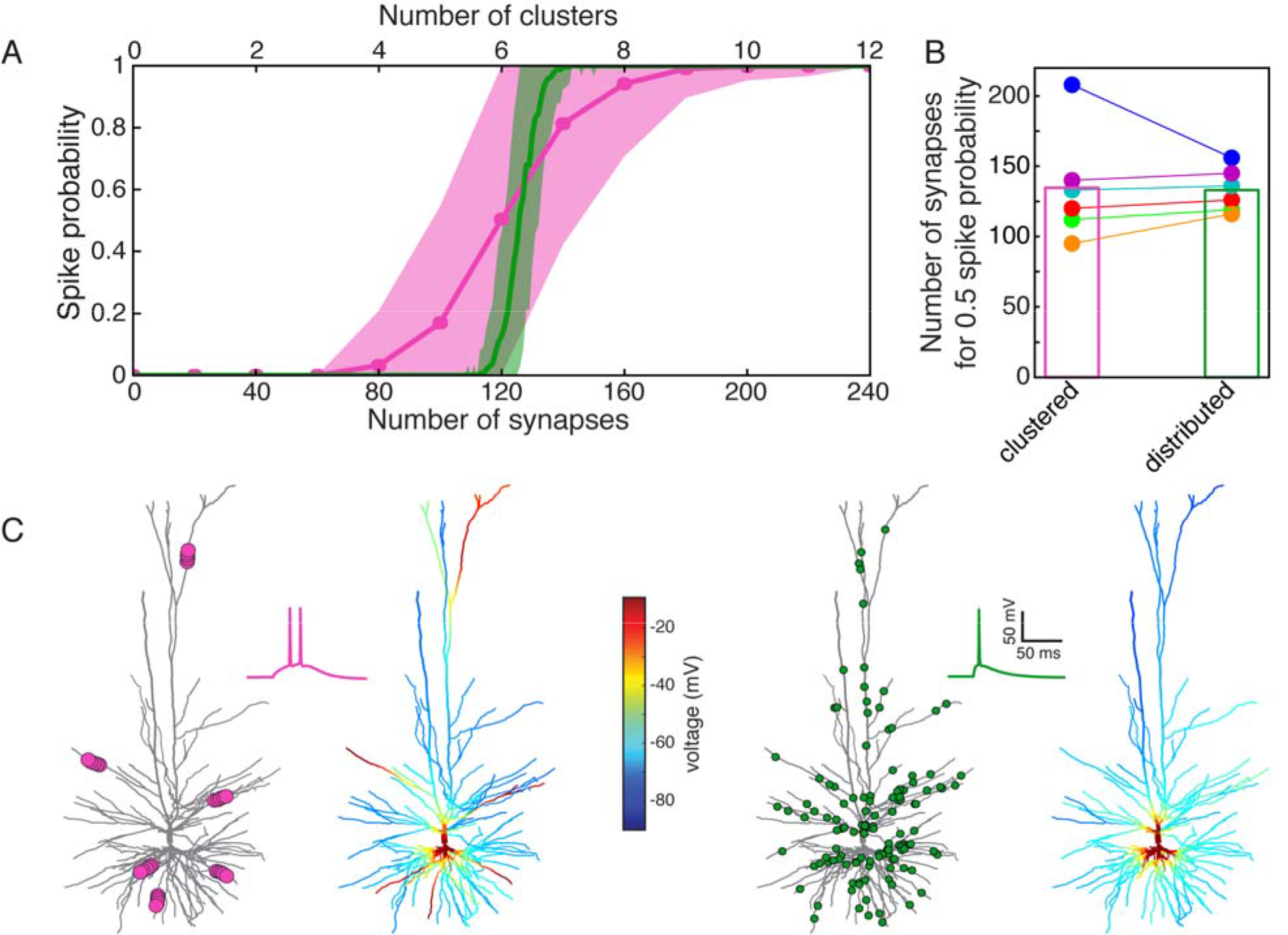
Model prediction of the number of excitatory HL2/L3 - HL2/L3 synapses required to generate a somatic Na^+^ spike. **(A)** Probability for a somatic spike presented as a function of the number of simultaneously activated synapses. Two cases are shown, randomly distributed synapses (green) and clustered synapses (pink), see **Methods**. **(B)** Number of synapses required to generate an AP with probability of 0.5 for the six HL2/L3 pyramidal cells modeled in Figure 7. Columns represent the mean value for the six models. The red case is for the neuron modeled in **(A)**; colors match the colors in Figure 7. **(C)** Example of a simulation for the clustered and the distributed cases for the model shown in **(A)**. Left, 6 clusters, 20 synapses each (pink dots at left tree; for illustration reasons only 4 synapses are shown per cluster), giving rise to local NMDA-spikes and a burst of two somatic Na^+^ spikes (pink trace). Corresponding color-coded spatial spread of voltage is depicted at the right tree. Right panel, 125 randomly distributed synapses (green dots) resulted in a single somatic spike (green trace) without dendritic NMDA spikes.

**Figure 9.**
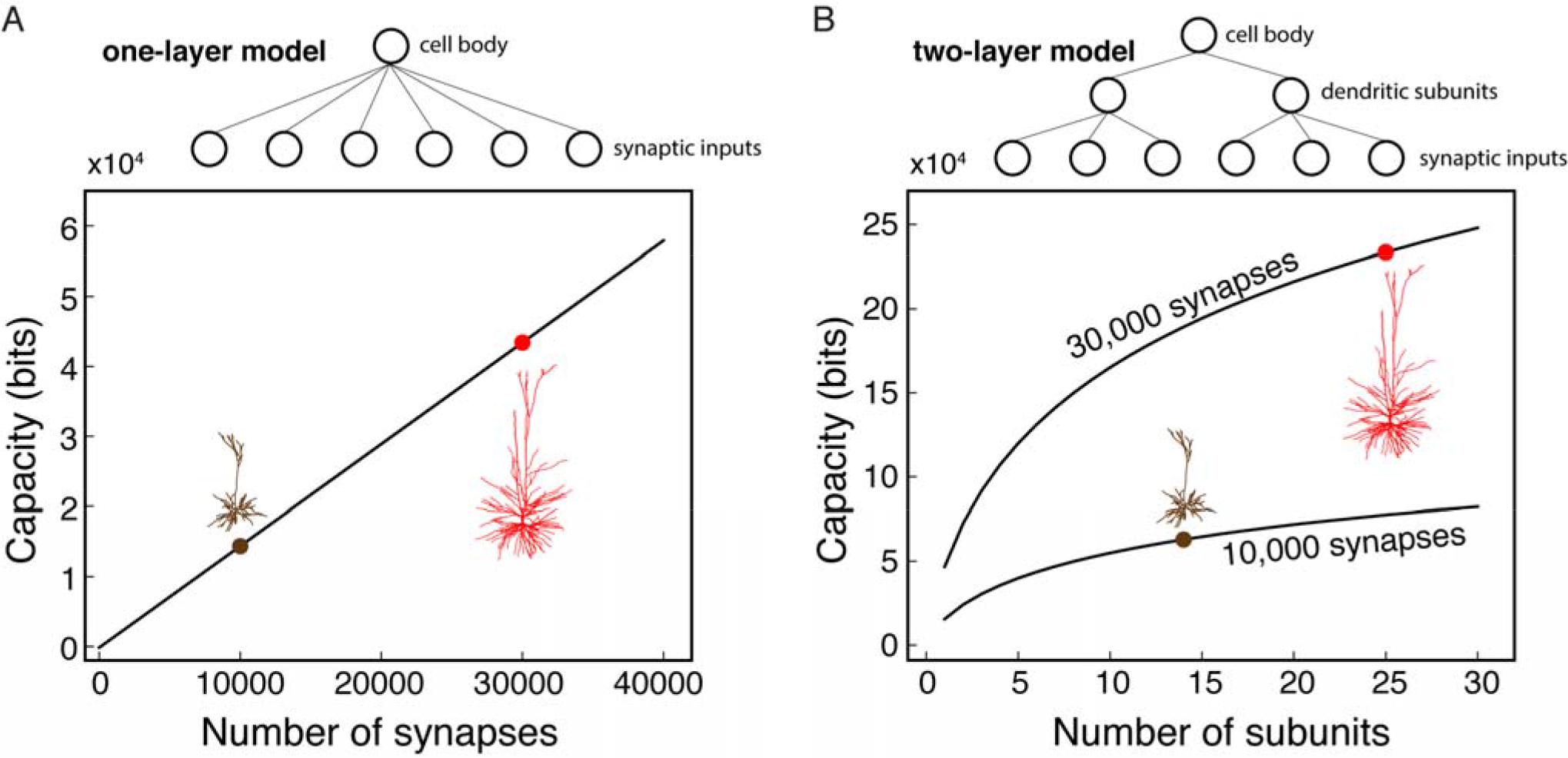
Human L2/L3 pyramidal neurons have larger storage capacity compared to rat. **(A)** Storage capacity as a function of the number of synaptic inputs when the neuron is considered as one-layer model. The inputs from all the synapses are summed directly at the cell body. The case of rodent L2/3 cell and HL2/L3 with 10,000 and 30,000 synapses is depicted, by black and red, respectively. **(B)** Storage capacity of the neuron when considered as a two-layer model; the capacity is shown as function of the number of non-linear dendritic subunits per neuron. The case of 14 subunits versus 25 subunits is shown for the rat and human PCs, respectively. Top and bottom curves are for 30,000 and 10,000 synaptic inputs respectively. The average number of contacts per connection was assumed to be five in both cases (d = s/5 for the parameters used in (Poirazi and Mel, 2001). Analysis of storage capacity is as in (Poirazi and Mel, 2001). Note that the capacity of human L2/L3 is almost four-folds that of the rodent.

## References

Alonso-Nanclares, L., Gonzalez-Soriano, J., Rodriguez, J. R., and DeFelipe, J. (2008). Gender differences in human cortical synaptic density. Proc. Natl. Acad. Sci. 105, 14615–14619.

Amunts, K., Ebell, C., Muller, J., Telefont, M., Knoll, A., and Lippert, T. (2016). The Human Brain Project: creating a European research infrastructure to decode the human brain. Neuron 92, 574–581.

Angelino, E., and Brenner, M. P. (2007). Excitability constraints on voltage-gated sodium channels. PLoS Comput. Biol. 3, 1751–60. doi:10.1371/journal.pcbi.0030177.

Araya, R., Vogels, T. P., and Yuste, R. (2014). Activity-dependent dendritic spine neck changes are correlated with synaptic strength. Proc. Natl. Acad. Sci. 111, E2895–E2904. doi:10.1073/pnas.1321869111.

Avoli, M., Louvel, J., Pumain, R., and Köhling, R. (2005). Cellular and molecular mechanisms of epilepsy in the human brain. Prog. Neurobiol. 77, 166–200.

Ballesteros-Yáñez, I., Benavides-Piccione, R., Elston, G. N., Yuste, R., and DeFelipe, J. (2006). Density and morphology of dendritic spines in mouse neocortex. Neuroscience 138, 403–409.

Benavides-Piccione, R., Arellano, J. I., and DeFelipe, J. (2005). Catecholaminergic innervation of pyramidal neurons in the human temporal cortex. Cereb. Cortex 15, 1584–1591.

Benavides-Piccione, R., Ballesteros-Yáñez, I., DeFelipe, J., and Yuste, R. (2002). Cortical area and species differences in dendritic spine morphology. J. Neurocytol. 31, 337–46. Available at: http://www.ncbi.nlm.nih.gov/pubmed/12815251 [Accessed August 5, 2013].

Benavides-Piccione, R., Fernaud-Espinosa, I., Robles, V., Yuste, R., and DeFelipe, J. (2013). Age-based comparison of human dendritic spine structure using complete three-dimensional reconstructions. Cereb. Cortex 23, 1798–1810. doi:10.1093/cercor/bhs154.

Blazquez-Llorca, L., Merchán-Pérez, Á., Rodríguez, J.-R., Gascón, J., and DeFelipe, J. (2013). FIB/SEM technology and Alzheimer’s disease: three-dimensional analysis of human cortical synapses. J. Alzheimer’s Dis. 34, 995–1013.

Bono, J., Wilmes, K. A., and Clopath, C. (2017). Modelling plasticity in dendrites: from single cells to networks. Curr. Opin. Neurobiol. 46, 136–141. doi:10.1016/j.conb.2017.08.013.

Brent, R. (1976). A new algorithm for minimizing a function of several variables without calculating derivatives. Algorithms Minimization without Deriv. Hall, Englewood Cliffs, NJ), 200–248.

Brodmann, K. (2007). Brodmann’s: Localisation in the Cerebral Cortex. Springer Science & Business Media.

Carnevale, N. T., and Hines, M. L. (2006). The NEURON book. Cambridge University Press.

DeFelipe, J. (2015). The anatomical problem posed by brain complexity and size: a potential solution. Front. Neuroanat. 9.

DeFelipe, J., Alonso-Nanclares, L., and Arellano, J. I. (2002). Microstructure of the neocortex: Comparative aspects. J. Neurocytol. 31, 299–316. doi:10.1023/A:1024130211265.

Deitcher, Y., Eyal, G., Kanari, L., Verhoog, M. B., Kahou, A., Antoine, G., et al. (2017). Comprehensive Morpho-Electrotonic Analysis Shows 2 Distinct Classes of L2 and L3 Pyramidal Neurons in Human Temporal Cortex. Cereb. Cortex, 1–17.

Druckmann, S., Banitt, Y., Gidon, A., Schürmann, F., Markram, H., and Segev, I. (2007). A novel multiple objective optimization framework for constraining conductance-based neuron models by experimental data. Front. Neurosci. 1, 7–18. doi:10.3389/neuro.01.1.1.001.2007.

Egger, R., Dercksen, V. J., Udvary, D., Hege, H.-C., and Oberlaender, M. (2014). Generation of dense statistical connectomes from sparse morphological data. Front. Neuroanat. 8, 129. doi:10.3389/fnana.2014.00129.

Elston, G. N., Benavides-Piccione, R., and DeFelipe, J. (2001). The pyramidal cell in cognition: a comparative study in human and monkey. J. Neurosci. 21, RC163. Available at: http://www.ncbi.nlm.nih.gov/pubmed/11511694.

Eyal, G., Mansvelder, H. D., de Kock, C. P. J., and Segev, I. (2014). Dendrites impact the encoding capabilities of the axon. J. Neurosci. 34, 8063–71. doi:10.1523/JNEUROSCI.5431-13.2014.

Eyal, G., Verhoog, M. B., Testa-Silva, G., Deitcher, Y., Lodder, J. C., Benavides-Piccione, R., et al. (2016). Unique membrane properties and enhanced signal processing in human neocortical neurons. Elife 5, e16553.

Farinella, M., Ruedt, D. T., Gleeson, P., Lanore, F., and Silver, R. A. (2014). Glutamate-bound NMDARs arising from in vivo-like network activity extend spatio-temporal integration in a L5 cortical pyramidal cell model. PLoS Comput. Biol. 10, e1003590. doi:10.1371/journal.pcbi.1003590.

Feldmeyer, D., Lübke, J., and Sakmann, B. (2006). Efficacy and connectivity of intracolumnar pairs of layer 2/3 pyramidal cells in the barrel cortex of juvenile rats. J. Physiol. 575, 583–602.

Feldmeyer, D., Lübke, J., Silver, R. A., and Sakmann, B. (2002). Synaptic connections between layer 4 spiny neurone-layer 2/3 pyramidal cell pairs in juvenile rat barrel cortex: physiology and anatomy of interlaminar signalling within a cortical column. J. Physiol. 538, 803–822.

Harnett, M. T., Makara, J. K., Spruston, N., Kath, W. L., and Magee, J. C. (2012). Synaptic amplification by dendritic spines enhances input cooperativity. Nature 491, 599. doi:10.1038/nature11554.

Hawrylycz, M., Anastassiou, C., Arkhipov, A., Berg, J., Buice, M., Cain, N., et al. (2016). Inferring cortical function in the mouse visual system through large-scale systems neuroscience. Proc. Natl. Acad. Sci. 113, 7337–7344.

Hay, E., Hill, S., Schürmann, F., Markram, H., and Segev, I. (2011). Models of neocortical layer 5b pyramidal cells capturing a wide range of dendritic and perisomatic active properties. PLoS Comput. Biol. 7, e1002107. doi:10.1371/journal.pcbi.1002107.

Hay, E., Schürmann, F., Markram, H., and Segev, I. (2013). Preserving axosomatic spiking features despite diverse dendritic morphology. J. Neurophysiol. 109, 2972–81. doi:10.1152/jn.00048.2013.

Hay, E., and Segev, I. (2014). Dendritic Excitability and Gain Control in Recurrent Cortical Microcircuits. Cereb. Cortex. doi:10.1093/cercor/bhu200.

Herz, A. V. M., Gollisch, T., Machens, C. K., and Jaeger, D. (2006). Modeling single-neuron dynamics and computations: a balance of detail and abstraction. Science (80-.). 314, 80–85.

Jadi, M. P., Behabadi, B. F., Poleg-Polsky, A., Schiller, J., and Mel, B. W. (2014). An augmented two-layer model captures nonlinear analog spatial integration effects in pyramidal neuron dendrites. Proc. IEEE 102, 782–798.

Jahr, C. E., and Stevens, C. F. (1990). Voltage dependence of NMDA-activated macroscopic conductances predicted by single-channel kinetics. J. Neurosci. 10, 3178–3182.

Kasthuri, N., Hayworth, K. J., Berger, D. R., Schalek, R. L., Conchello, J. A., Knowles-Barley, S., et al. (2015). Saturated Reconstruction of a Volume of Neocortex. Cell 162, 648–661. doi:10.1016/j.cell.2015.06.054.

Koch, C., and Jones, A. (2016). Big science, team science, and open science for neuroscience. Neuron 92, 612–616.

Koch, C., Poggio, T., and Torres, V. (1982). Retinal ganglion cells: a functional interpretation of dendritic morphology. Philos. Trans. R. Soc. B Biol. Sci. 298, 227–263.

Koch, C., and Segev, I. (2000). The role of single neurons in information processing. Nat. Neurosci. 3 Suppl, 1171–1177.

Köhling, R., and Avoli, M. (2006). Methodological approaches to exploring epileptic disorders in the human brain in vitro. J. Neurosci. Methods 155, 1–19. doi:10.1016/j.jneumeth.2006.04.009.

Kole, M. H., and Stuart, G. J. (2012). Signal processing in the axon initial segment. Neuron 73, 235–247. doi:10.1016/j.neuron.2012.01.007.

Kwon, T., Sakamoto, M., Peterka, D. S., and Yuste, R. (2017). Attenuation of synaptic potentials in dendritic spines. Cell Rep. 20, 1100–1110.

Larkum, M. E., Nevian, T., Sandler, M., Polsky, A., and Schiller, J. (2009). Synaptic integration in tuft dendrites of layer 5 pyramidal neurons: a new unifying principle. Science 325, 756–60. doi:10.1126/science.1171958.

Larkum, M. E., Zhu, J. J., and Sakmann, B. (1999). A new cellular mechanism for coupling inputs arriving at different cortical layers. Nature 398, 338–341.

Lavzin, M., Rapoport, S., Polsky, A., Garion, L., and Schiller, J. (2012). Nonlinear dendritic processing determines angular tuning of barrel cortex neurons in vivo. Nature 490, 397–401. doi:10.1038/nature11451.

Magee, J. C., and Cook, E. P. (2000). Somatic EPSP amplitude is independent of synapse location in hippocampal pyramidal neurons. Nat. Neurosci. 3, 895.

Major, G., Larkum, M. E., and Schiller, J. (2013). Active properties of neocortical pyramidal neuron dendrites. Annu. Rev. Neurosci. 36, 1–24. doi:10.1146/annurev-neuro-062111-150343.

Markram, H., Lübke, J., Frotscher, M., Roth, A., and Sakmann, B. (1997a). Physiology and anatomy of synaptic connections between thick tufted pyramidal neurones in the developing rat neocortex. J. Physiol. 500, 409–440.

Markram, H., Lübke, J., Frotscher, M., and Sakmann, B. (1997b). Regulation of synaptic efficacy by coincidence of postsynaptic APs and EPSPs. Science (80-.). 275, 213–215.

Markram, H., Muller, E., Ramaswamy, S., Reimann, M. W., Abdellah, M., Sanchez, C. A., et al. (2015). Reconstruction and Simulation of Neocortical Microcircuitry. Cell 163, 456–492.

Martin, C. L., and Chun, M. (2016). The BRAIN initiative: Building, strengthening, and sustaining. Neuron 92, 570–573.

McCulloch, W. S., and Pitts, W. (1943). A logical calculus of the ideas immanent in nervous activity. Bull. Math. Biophys. 5, 115–133.

Mel, B. W. (1992). NMDA-based pattern discrimination in a modeled cortical neuron. Neural Comput. 4, 502–517.

Mel, B. W., Schiller, J., and Poirazi, P. (2017). Synaptic plasticity in dendrites: complications and coping strategies. Curr. Opin. Neurobiol. 43, 177–186.

Mohan, H., Verhoog, M. B., Doreswamy, K. K., Eyal, G., Aardse, R., Lodder, B. N., et al. (2015). Dendritic and axonal architecture of individual pyramidal neurons across layers of adult human neocortex. Cereb. Cortex 25, 4839–4853. Available at: http://cercor.oxfordjournals.org/content/25/12/4839.short.

Molnár, G., Rózsa, M., Baka, J., Holderith, N., Barzó, P., Nusser, Z., et al. (2016). Human pyramidal to interneuron synapses are mediated by multi-vesicular release and multiple docked vesicles. Elife 5, e18167.

Palmer, L. M., Shai, A. S., Reeve, J. E., Anderson, H. L., Paulsen, O., and Larkum, M. E. (2014). NMDA spikes enhance action potential generation during sensory input. Nat. Neurosci. 17, 383–90. doi:10.1038/nn.3646.

Palmer, L. M., and Stuart, G. J. (2009). Membrane potential changes in dendritic spines during action potentials and synaptic input. J. Neurosci. 29, 6897–6903.

Poirazi, P., Brannon, T., and Mel, B. W. (2003a). Arithmetic of subthreshold synaptic summation in a model CA1 pyramidal cell. Neuron 37, 977–987.

Poirazi, P., Brannon, T., and Mel, B. W. (2003b). Pyramidal neuron as two-layer neural network. Neuron 37, 989–99. Available at: http://www.ncbi.nlm.nih.gov/pubmed/12670427 [Accessed August 9, 2015].

Poirazi, P., and Mel, B. W. (2001). Impact of Active Dendrites and Structural Plasticity on the Memory Capacity of Neural Tissue. Neuron 29, 779–796. doi:10.1016/S0896-6273(01)00252-5.

Polsky, A., Mel, B. W., and Schiller, J. (2004). Computational subunits in thin dendrites of pyramidal cells. Nat. Neurosci. 7, 621–627.

Poo, M., Du, J., Ip, N. Y., Xiong, Z.-Q., Xu, B., and Tan, T. (2016). China Brain Project: basic neuroscience, brain diseases, and brain-inspired computing. Neuron 92, 591–596.

Popovic, M. A., Carnevale, N., Rozsa, B., and Zecevic, D. (2015). Electrical behaviour of dendritic spines as revealed by voltage imaging. Nat. Commun. 6.

Rall, W. (1959). Branching dendritic trees and motoneuron membrane resistivity. Exp. Neurol. 1, 491–527. doi:10.1016/0014-4886(59)90046-9.

Rall, W. (1964). Theoretical significance of dendritic trees for neuronal input-output relations Neural Theory Model., 73–97.

Rall, W. (1967). Distinguishing theoretical synaptic potentials computed for different soma-dendritic distributions of synaptic input. J. Neurophysiol. 30, 1138–68. Available at: http://www.ncbi.nlm.nih.gov/pubmed/6055351 [Accessed August 4, 2013].

Rall, W. (1969). Time constants and electrotonic length of membrane cylinders and neurons. Biophys. J. 9, 1483–508. doi:10.1016/S0006-3495(69)86467-2.

Rall, W., Burke, R. E., Smith, T. G., Nelson, P. G., and Frank, K. (1967). Dendritic location of synapses and possible mechanisms for the monosynaptic EPSP in motoneurons. J. Neurophysiol 30, 884–915.

Ranjan, R., Khazen, G., Gambazzi, L., Ramaswamy, S., Hill, S. L., Schürmann, F., et al. (2011). Channelpedia: an integrative and interactive database for ion channels. Front. Neuroinform. 5, 36. doi:10.3389/fninf.2011.00036.

Rapp, M., Yarom, Y., and Segev, I. (1992). The impact of parallel fiber background activity on the cable properties of cerebellar Purkinje cells. Neural Comput. 4, 518–533.

Rhodes, P. (2006). The properties and implications of NMDA spikes in neocortical pyramidal cells. J. Neurosci. 26, 6704–6715.

del Río, M. R., and DeFelipe, J. (1994). A study of SMI 32-stained pyramidal cells, parvalbumin-immunoreactive chandelier cells, and presumptive thalamocortical axons in the human temproal neocortex. J. Comp. Neurol. 342, 389–408.

Sarid, L., Bruno, R., Sakmann, B., Segev, I., and Feldmeyer, D. (2007). Modeling a layer 4-to-layer 2/3 module of a single column in rat neocortex: interweaving in vitro and in vivo experimental observations. Proc. Natl. Acad. Sci. U. S. A. 104, 16353–8. doi:10.1073/pnas.0707853104.

Sarid, L., Feldmeyer, D., Gidon, A., Sakmann, B., and Segev, I. (2013). Contribution of Intracolumnar Layer 2/3-to-Layer 2/3 Excitatory Connections in Shaping the Response to Whisker Deflection in Rat Barrel Cortex. Cereb. Cortex. doi:10.1093/cercor/bht268.

Schiller, J., Major, G., Koester, H. J., and Schiller, Y. (2000). NMDA spikes in basal dendrites of cortical pyramidal neurons. Nature 404, 285–289.

Schmidt-Hieber, C., Toleikyte, G., Aitchison, L., Roth, A., Clark, B. A., Branco, T., et al. (2017). Active dendritic integration as a mechanism for robust and precise grid cell firing. Nat. Neurosci.

Segev, I., Friedman, A., White, E. L., and Gutnick, M. J. (1995). Electrical consequences of spine dimensions in a model of a cortical spiny stellate cell completely reconstructed from serial thin sections. J. Comput. Neurosci. 2, 117–130. doi:10.1007/BF00961883.

Shalev-Shwartz, S., and Ben-David, S. (2014). Understanding machine learning: From theory to algorithms. Cambridge university press.

Shen, G. Y., Chen, W. R., Midtgaard, J., Shepherd, G. M., and Hines, M. L. (1999). Computational analysis of action potential initiation in mitral cell soma and dendrites based on dual patch recordings. J. Neurophysiol. 82, 3006–3020.

Shimizu, E., Tang, Y.-P., Rampon, C., and Tsien, J. Z. (2000). NMDA receptor-dependent synaptic reinforcement as a crucial process for memory consolidation. Science (80-.). 290, 1170–1174.

Smith, S. L., Smith, I. T., Branco, T., and Häusser, M. (2013). Dendritic spikes enhance stimulus selectivity in cortical neurons in vivo. Nature 503, 115–20. doi:10.1038/nature12600.

Spruston, N. (2008). Pyramidal neurons: dendritic structure and synaptic integration. Nat:. Rev. Neurosci. 9, 206–221.

Stuart, G. J., and Sakmann, B. (1994). Active propagation of somatic action potentials into neocortical pyramidal cell dendrites. Nature 367, 69–72. doi:10.1038/367069a0.

Stuart, G., Spruston, N., and Häusser, M. (2016). Dendrites. Oxford University Press.

Svoboda, K., Tank, D. W., and Denk, W. (1996). Direct measurement of coupling between dendritic spines and shafts. Science 272, 716–9. Available at: http://www.ncbi.nlm.nih.gov/pubmed/8614831 [Accessed April 5, 2015].

Szabadics, J., Varga, C., Molnár, G., Oláh, S., Barzó, P., and Tamás, G. (2006). Excitatory effect of GABAergic axo-axonic cells in cortical microcircuits. Science 311, 233–5. doi:10.1126/science.1121325.

Testa-Silva, G., Verhoog, M. B., Goriounova, N. a, Loebel, A., Hjorth, J., Baayen, J. C., et al. (2010). Human synapses show a wide temporal window for spike-timing-dependent plasticity. Front. Synaptic Neurosci. 2, 12. doi:10.3389/fnsyn.2010.00012.

Testa-Silva, G., Verhoog, M. B., Linaro, D., de Kock, C. P. J., Baayen, J. C., Meredith, R. M., et al. (2014). High bandwidth synaptic communication and frequency tracking in human neocortex. PLoS Biol. 12, e1002007. doi:10.1371/journal.pbio.1002007.

Tian, C., Wang, K., Ke, W., Guo, H., and Shu, Y. (2014). Molecular identity of axonal sodium channels in human cortical pyramidal cells. Front. Cell. Neurosci. 8, 1–16. doi:10.3389/fncel.2014.00297.

Tønnesen, J., Katona, G., Rózsa, B., and Nägerl, U. V. (2014). Spine neck plasticity regulates compartmentalization of synapses. Nat. Neurosci. 17, 678–685.

Varga, C., Tamas, G., Barzo, P., Olah, S., and Somogyi, P. (2015). Molecular and Electrophysiological Characterization of GABAergic interneurons expressing the transcription factor COUP-TFII in the adult human temporal cortex. Cereb. Cortex 25, 4430–4449.

Verhoog, M. B., Goriounova, N. A., Obermayer, J., Stroeder, J., Hjorth, J. J. J., Testa-Silva, G., et al. (2013). Mechanisms underlying the rules for associative plasticity at adult human neocortical synapses. J. Neurosci. 33, 17197–208. doi:10.1523/JNEUROSCI.3158-13.2013.

Wuarin, J. P., Peacock, W. J., and Dudek, F. E. (1992). Single-electrode voltage-clamp analysis of the N-methyl-D-aspartate component of synaptic responses in neocortical slices from children with intractable epilepsy. J Neurophysiol 67, 84–93. Available at: http://jn.physiology.org/content/67/1/84.full-text.pdf+html [Accessed July 27, 2015].

